# Spatial organization of fibroblast subpopulations and niches across acute repair and chronic fibrotic remodeling of the human myocardium

**DOI:** 10.64898/2026.10.06.756898

**Authors:** Alexandru Micu, Alexandru-Emil Matei, Yi-Nan Li, Ann-Christin Pecher, Tim Filla, Peter-Martin Bruch, Sascha Dietrich, Jörg Henes, Elric Zweck, Markus Eckstein, Karin Klingel, Jörg H.W. Distler, Andrea-Hermina Györfi

**Affiliations:** Department of Rheumatology, University Hospital Düsseldorf, Medical Faculty of Heinrich-Heine University; Düsseldorf, Germany; Hiller Research Center, University Hospital Düsseldorf, Medical Faculty of Heinrich-Heine University; Düsseldorf, Germany; Fraunhofer Institute for Translational Medicine and Pharmacology ITMP, and Fraunhofer Cluster of Excellence for Immune Mediated Diseases CIMD, Frankfurt am Main, Germany; Spatial & Functional Screening Core Facility, Medical Faculty, Heinrich Heine University Düsseldorf, Düsseldorf, Germany; Department of Internal Medicine II, Hematology, Oncology, Clinical Immunology, and Rheumatology, University Hospital Tübingen, Tübingen, Germany; Department of Hematology, Oncology and Clinical Immunology, Medical Faculty and University Hospital Düsseldorf, Düsseldorf, Germany; Department of Cardiology, Pulmonology and Vascular Medicine, University Hospital Duesseldorf, Duesseldorf, Germany; Institute of Pathology, Friedrich-Alexander-Universität Erlangen-Nürnberg, Universitätsklinikum Erlangen, and Deutsches Zentrum für Immuntherapie (DZI), Erlangen, Germany; Cardiopathology, Institute for Pathology, Eberhard-Karls-Universität Tübingen, Tübingen, Germany

## Abstract

Myocardial fibrosis is a common end point of acute and chronic cardiovascular injury, yet the spatially organized fibroblast (Fb) and immune-cell states that sustain fibrotic remodeling across etiologies remain incompletely defined. We applied cyclic *in situ* hybridization (cISH)-based spatial transcriptomics to myocardial biopsies from patients with acute myocardial infarction (AMI), ischemic cardiomyopathy (ICM), hypertrophic cardiomyopathy (HCM), postviral myocardial fibrosis (PVMF), and systemic sclerosis associated primary heart involvement (SSc-pHI). CODEX-based protein imaging provided orthogonal support for cell subpopulations and cellular neighborhoods. Spatially informed clustering identified six Fb subpopulations: COL1A1^hi^/POSTN^hi^, NOTCH3^+^, TIMP1^hi^, GSN^hi^/DCN^hi^, AP1^hi^ Fbs, and myofibroblasts. COL1A1^hi^/POSTN^hi^ and NOTCH3^+^ Fbs were enriched in AMI, whereas GSN^hi^/DCN^hi^ Fbs and myofibroblasts predominated in chronic fibrotic remodeling. These subpopulations displayed distinct extracellular matrix, inflammatory, and vasoregulatory programs. Infarction-associated reparative niches were enriched in COL1A1^hi^/POSTN^hi^ or NOTCH3^+^ Fbs and SPP1^+^ or APOE^+^ macrophages (MФ), whereas chronic fibroinflammatory niches were enriched in GSN^hi^/DCN^hi^ or AP1^hi^ Fbs or myofibroblasts with C1Q^+^ MФ and plasma cells. The GSN^hi^/DCN^hi^ Fb niche marked areas of subendocardial fibrosis, whereas the AP1^hi^ Fb niche localized to perivascular fibrosis. In SSc-pHI, the actively fibrogenic COL1A1^hi^/POSTN^hi^ Fb state characteristic of infarct scar formation was present at higher frequency than in other chronic entities and was individually associated with cardiac dysfunction, whereas the chronic GSN^hi^/DCN^hi^ Fb niche was associated with the extent of interstitial fibrosis and anti-topoisomerase I autoantibodies. These data show that human myocardial fibrosis is organized into spatially distinct Fb subpopulations that assemble into context-specific fibroinflammatory niches across chronic fibrotic remodeling and repair after AMI.

**What Is New?:**

- Spatially resolved single-cell transcriptomics of myocardial biopsies from acute myocardial infarction, ischemic and hypertrophic cardiomyopathy, postviral myocardial fibrosis, and systemic sclerosis-associated primary heart involvement (SSc-pHI) resolves six fibroblast subpopulations assembling into context-specific fibroinflammatory niches.
- Infarct scar formation and chronic fibrotic remodeling are organized by distinct fibroblast-macrophage circuits: COL1A1^hi^/POSTN^hi^ and NOTCH3^+^ fibroblasts with SPP1^+^ macrophages in infarct scar, versus GSN^hi^/DCN^hi^ and AP1^hi^ fibroblasts with C1Q^+^ macrophages and plasma cells in chronic disease. Subendocardial fibrosis maps onto GSN^hi^/DCN^hi^ and perivascular fibrosis onto AP1hi neighborhoods.
- SSc-pHI is distinguished by increased actively fibrogenic COL1A1^hi^/POSTN^hi^ fibroblasts, transient during infarct scar formation, indicating ongoing fibrogenesis.

**What Are the Clinical Implications?:**

- Spatial profiling links histological patterns of myocardial fibrosis to distinct fibroblast and immune-cell environments, providing a basis for investigating how these cellular relationships contribute to myocardial remodeling.
- In SSc-pHI, infarct scar-associated COL1A1^hi^/POSTN^hi^ fibroblasts occur alongside GSN^hi^/DCN^hi^ fibroblasts. These findings provide a basis for functional and longitudinal studies to define their contributions to myocardial remodeling and evaluate their potential as markers of disease activity or therapeutic targets.
- Prospective studies integrating these tissue findings with cardiac imaging and circulating biomarkers should establish whether the observed cellular patterns provide additional information about disease progression or treatment response.

## Introduction

Myocardial fibrosis is a shared pathological end point of different cardiovascular injuries, encompassing reparative scar formation after acute injury and persistent remodeling in chronic disease. Human cardiac atlases have established fibroblast heterogeneity, including regionally specialized states in non-diseased myocardium, and identified functionally distinct macrophage subsets [1,2]. Understanding how these populations assemble into local tissue niches is essential to defining the cellular organization of different forms of myocardial remodeling.

Human single-cell and spatial studies have mapped activated fibroblast states and immune responses after acute myocardial infarction (AMI), in ischemic cardiomyopathy (ICM), and in hypertrophic cardiomyopathy (HCM) [3–6]. Recent work has further linked fibroblast activation and vascular inflammatory programs to regional and histological features of failing hearts [7,8]. Building on these advances, a central unresolved question is which fibroblast-immune niches are shared across chronic fibrotic etiologies, which vary by etiology, and which distinguish chronic remodeling from acute infarct repair. Addressing this question requires comparative assessment of cell states, local immune partners, and spatial organization within a common analytical framework.

This gap is particularly relevant to postviral myocardial fibrosis (PVMF) and systemic sclerosis-associated primary heart involvement (SSc-pHI). In postviral myocardial disease, immune-mediated mechanisms have been linked to persistent dysfunction and fibrotic transcriptional responses [9–11]. In SSc-pHI, histopathology and molecular imaging document myocardial fibrosis, inflammation, and fibroblast activation associated with adverse cardiac phenotypes [12–14]. Yet the fibroblast-immune niches associated with distinct histological patterns of fibrosis, and their relationship to cardiac manifestations in SSc-pHI, remain to be defined.

To address these questions, we applied cyclic in situ hybridization-based spatial transcriptomics to myocardial biopsies from 38 patients with AMI, ICM, HCM, PVMF, or SSc-pHI. We aimed to define fibroblast and immune-cell subpopulations in their tissue context, compare their niches and inferred interaction networks across acute repair and chronic fibrotic etiologies, and relate their spatial organization to histological fibrosis patterns. Matched-donor CODEX multiplex protein imaging was used to validate selected subpopulations and neighborhood components at the protein level. In SSc-pHI, we further sought to determine whether these spatial features were associated with myocardial fibrosis burden and cardiac manifestations.

## Methods

### Patients and endomyocardial samples

We included a total of 38 patients who underwent myocardial biopsy for diagnostic or therapeutic purposes: 13 SSc-pHI, eight PVMF, nine ICM, five HCM, and three AMI patients (Figure 1A).

**Figure 1:**
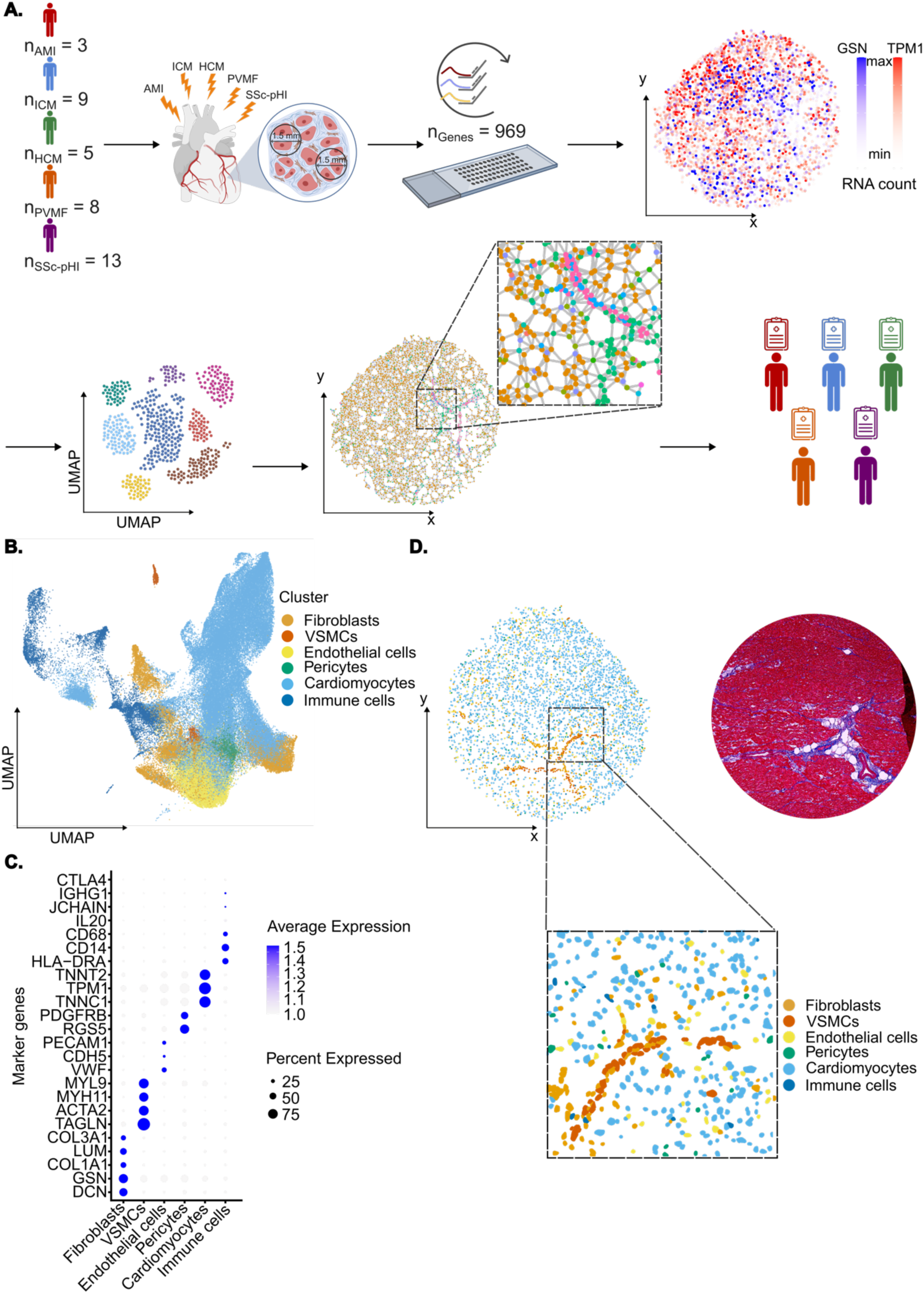
Spatial transcriptomic profiling and cellular annotation of myocardial biopsies. **(A)** Schematic overview of the study design and analytical workflow. Myocardial biopsies were collected from patients with acute myocardial infarction (AMI, n = 3), ischemic cardiomyopathy (ICM, n = 9), hypertrophic cardiomyopathy (HCM, n = 5), post-viral myocardial fibrosis (PVMF, n = 8), and systemic sclerosis primary heart involvement (SSc-pHI, n = 13). Two 1.5-mm tissue punches were obtained from each biopsy and subjected to cyclic in situ hybridization. Spatially resolved transcriptomic profiles were generated at single-cell resolution across 969 genes. Representative spatial expression patterns of GSN and TPM1 are shown. Cells were clustered and annotated based on transcriptional profiles, followed by spatial neighborhood and cell-cell interaction analyses. Spatial and cellular features were subsequently correlated with clinical parameters. **(B)** Uniform manifold approximation and projection (UMAP) embedding of all cells showing the major annotated cell populations. **(C)** Dot plot of canonical marker genes used for cell-type annotation. Dot size represents the proportion of cells expressing a given gene, and color intensity indicates average expression level. **(D)** Representative Masson’s trichrome-stained myocardial section and corresponding spatial map of annotated cells, illustrating the localization of major cell populations within the tissue microenvironment. Insets show higher-magnification views of the spatial organization of annotated cells.

Biopsy material was retrieved via right-heart catheterization from the septal wall of the right ventricle in SSc-pHI, while PVMF, HCM, ICM and AMI patients’ biopsies derived from myocardial tissue retrieved from the left ventricle. FFPE material was further processed for analysis. From each individual biopsy, two circular regions with a diameter of 1.5 mm (corresponding to an area of 1.8 mm² per region) were selected for cISH. For AMI samples, Masson’s trichrome staining was performed on the immediately preceding serial section, and regions displaying histological features compatible with myocardial infarction were selected as the sites for TMA core punching.

Detailed descriptions of the materials and methods are provided in the Supplemental Methods section.

## Results

### Study design and identification of major cell types

We recovered a total of 109,632 cells (AMI: 14,242 cells; ICM: 29,535 cells HCM: 24,950 cells; PVMF: 14,783 cells; SSc-pHI: 26,122 cells). These single cells included 11,672 Fb (AMI: 2,178 cells; ICM: 2,118 cells HCM: 3,255 cells; PVMF: 1,246 cells; SSc-pHI: 2,875 cells) identified by expression of DCN [1], GSN [1], COL1A1 [3,5], LUM [39] and COL3A1 [3,5], and 7,794 immune cells identified by expression of the pan-leukocyte marker PTPRC/CD45 and lineage markers including CD14, CD68, HLA-DRA, JCHAIN, or IGHG1 (Figure 1C). We further identified other major stromal and parenchymal cell populations, including endothelial cells (ECs; PECAM1, CDH5, VWF), pericytes (RGS5, PDGFRB), vascular smooth muscle cells (VSMCs; TAGLN, ACTA2, MYH11, MYL9) (Figure 1C), and cardiomyocytes (TNNT2, TPM1, TNNC1). These populations showed the expected spatial distribution within myocardial tissue, supporting the validity of the cell annotations (Figure 1D).

### Spatially informed Fb subpopulations in human myocardial fibrosis

Using Building Aggregates with a Neighborhood Kernel and Spatial Yardstick (BANKSY) [23], which incorporates both intrinsic gene expression and neighborhood-derived transcriptional information, we identified six spatially informed Fb subpopulations, which were annotated according to their dominant marker combinations. These included previously described Fb subpopulations such as COL1A1^hi^/POSTN^hi^ Fbs, GSN^hi^/DCN^hi^ Fbs, AP1^hi^ Fbs and myofibroblasts, as well as TIMP1^hi^ and NOTCH3^+^ Fbs, which have not been described as discrete states in dissociation-based atlases. Notably, AP1^hi^ (which showed a primarily perivascular distribution) and TIMP1^hi^ Fbs were resolved only by spatially informed clustering, underscoring the importance of tissue context for identifying niche-dependent Fb states (Figure 2A–C). To place the six spatially informed Fb populations in the context of existing cardiac atlases, we performed scANVI label transfer using genes shared between our cISH panel and four published sc/snRNASeq datasets [1,3,29,30]. COL1A1^hi^/POSTN^hi^ Fbs showed prominent correspondence with activated fibroblast and myofibroblast reference populations. GSN^hi^/DCN^hi^ Fbs mapped across several reference fibroblasts subtypes, while AP1^hi^, TIMP1^hi^, and NOTCH3^+^ Fbs showed partially shared reference assignments. These comparisons related the BANKSY-defined populations to previously described fibroblast transcriptional states, with individual reference categories encompassing multiple spatially informed populations (Supplementary Figure 1).

**Figure 2:**
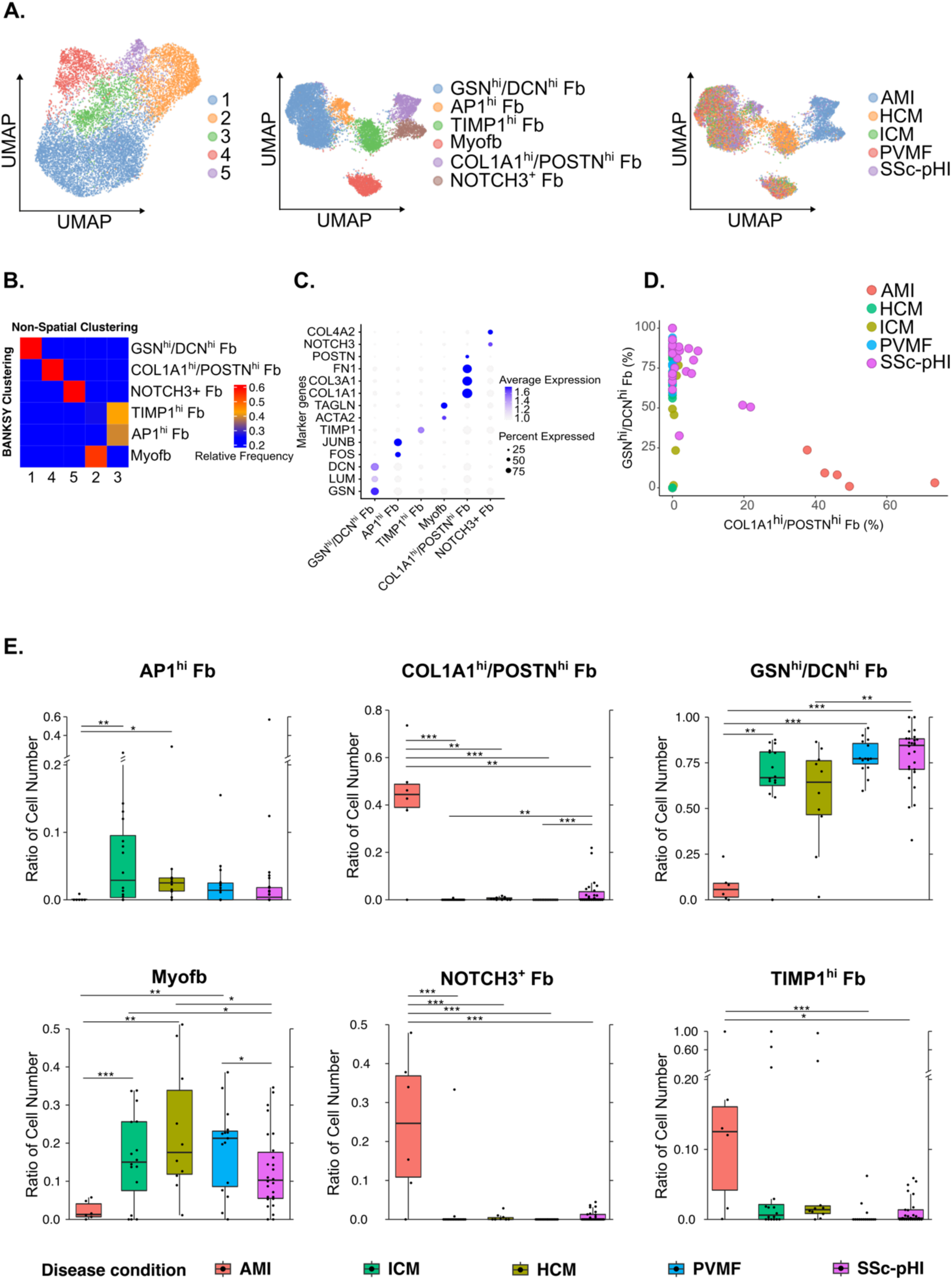
Spatially informed clustering identifies distinct fibroblast states across human myocardial fibrotic disease entities. **(A)** Fibroblast populations were first resolved by non-spatial clustering and visualized by UMAP. Spatially informed clustering was then performed using BANKSY, revealing fibroblast subclusters and their distribution across disease entities. **(B)** Heatmap comparing non-spatial and spatial clustering showing refinement of the fibroblast compartment by spatial analysis, including separation of non-spatial cluster 3 into AP1^hi^ and TIMP1^hi^ fibroblast states. **(C)** Dot plot of marker genes defining the spatially informed fibroblast subclusters. Dot size indicates the proportion of cells expressing each gene, and color intensity represents average expression level. **(D)** Scatter plot showing the proportions of COL1A1^hi^/POSTN^hi^ fibroblasts and GSN^hi^/DCN^hi^ fibroblasts, colored by disease condition. **(E)** Box plots showing the distribution of fibroblast subclusters across disease conditions. Each dot represents an individual sample. Statistical significance (Kruskal-Wallis test) is indicated as *p < 0.05, **p < 0.01, and ***p < 0.001.

### The distribution of Fb subpopulations differs across acute repair and chronic fibrotic remodeling contexts

COL1A1^hi^/POSTN^hi^ and NOTCH3^+^ Fbs were particularly enriched in AMI, consistent with repair in response to acute ischemic injury, whereas GSN^hi^/DCN^hi^ Fbs and myofibroblasts were enriched in chronic fibrotic remodeling relative to AMI (Figure 2E). Among the chronic fibrotic conditions, COL1A1^hi^/POSTN^hi^ Fbs were more abundant in SSc-pHI than in ICM, HCM, and PVMF. GSN^hi^/DCN^hi^ Fbs were also most frequent in SSc-pHI. SSc-pHI was thus the only chronic entity in which both the infarct-associated COL1A1^hi^/POSTN^hi^ Fbs and the chronic-remodeling-associated GSN^hi^/DCN^hi^ Fbs were enriched (Figure 2D-E). AP1^hi^ Fbs were numerically increased in chronic fibrotic remodeling, particularly in ICM, whereas TIMP1^hi^ Fbs were enriched in acute repair (Figure 2E).

### Spatially informed Fb subpopulations are functionally distinct and exhibit shared and disease-specific transcriptional programs across acute repair and chronic myocardial fibrosis

We next assessed the functional activation states of the Fb subpopulations. Matrix-related scoring using Matrisome-derived gene sets showed that COL1A1^hi^/POSTN^hi^ Fbs had the highest overall extracellular matrix (ECM) score, whereas GSN^hi^/DCN^hi^ and AP1^hi^ Fbs showed the highest glycoprotein and proteoglycan scores, consistent with distinct contributions to ECM composition. The activity of the transforming growth factor-β (TGFβ) pathway as a core pathway of fibrotic remodeling was highest in COL1A1^hi^/POSTN^hi^ Fbs (Figure 3A).

**Figure 3:**
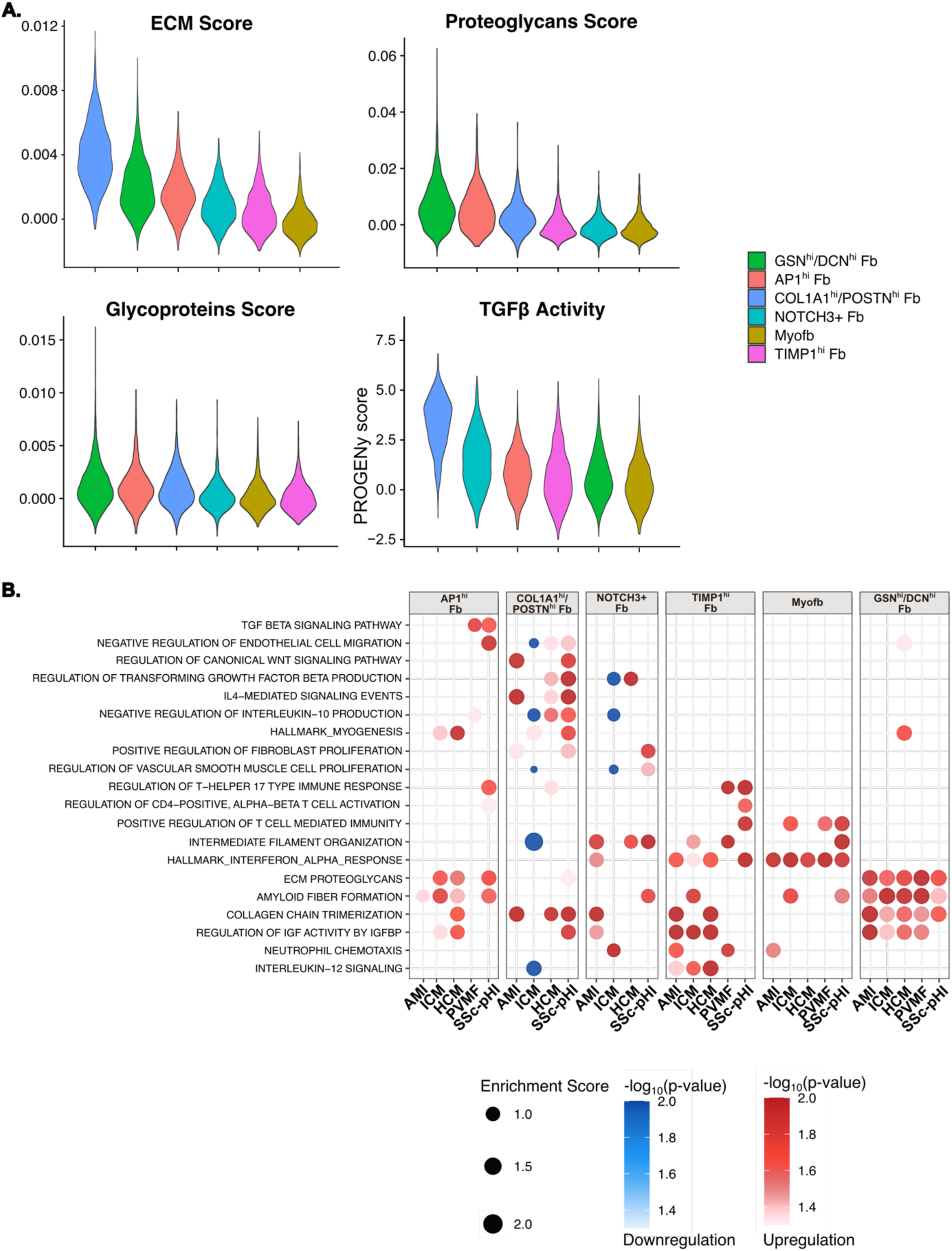
Functional programs of fibroblast subpopulations across disease conditions. **(A)** Violin plots showing ECM, proteoglycan, and glycoprotein scores based on gene sets curated in the Human Matrisome Project, together with TGF-β pathway activity inferred using PROGENy, across the distinct fibroblast subclusters. **(B)** Dot plot of selected pathway enrichment scores across fibroblast populations under different disease conditions. Enrichment analysis was performed using fgsea on pseudo-bulk expression matrices generated by averaging expression within each fibroblast subpopulation for each disease condition and implemented through the decoupleR R package. Pathways with p < 0.05 were retained. Statistical significance was assessed by comparing estimated marginal means from linear mixed-effects models including donor-specific random intercepts. Dot size indicates enrichment score, and color intensity represents normalized−log10(p) values for up- and downregulation.

Functional enrichment analysis further supported functional specialization of individual Fb subpopulations. COL1A1^hi^/POSTN^hi^ Fbs were enriched for ECM- and collagen-related programs together with canonical profibrotic pathways, including TGFβ, WNT and IL-4 signaling. TIMP1^hi^ Fbs showed increased IL-12-associated signaling; in SSc-pHI, they additionally exhibited signatures linked to T cell-mediated immunity. NOTCH3^+^ Fbs were enriched for programs related to Fb and VSMC proliferation in SSc-pHI. In accordance with their perivascular localization, AP1^hi^ Fbs showed activation of programs related to lymphocyte and endothelial-cell activation. Among all Fb subpopulations, AP1^hi^ Fbs demonstrated the strongest enrichment pathways involved in endothelial-cell regulation, most prominently shown in SSc-pHI. Of note, GSN^hi^/DCN^hi^ and AP1^hi^ Fbs additionally showed enrichment of myogenesis-related programs in HCM consistent with the hypertrophic myocardial phenotype. As expected, myofibroblasts were enriched for pathways related to cytoskeletal organization, including Rho GTPase-mediated regulation. Myofibroblasts also showed enrichment of immune-regulatory programs, including genes involved in T cell cytokine production and responses to interferon-*a*. Together, these data demonstrate that BANKSY-defined Fb subpopulations differ not only in spatial distribution but also in functional specialization across disease contexts (Figure 3B).

### Trajectory analysis identifies differentiation route from GSN^hi^/DCN^hi^ fibroblasts toward profibrotic fibroblasts

Monocle v3 pseudotime analysis was rooted in the GSN^hi^/DCN^hi^ fibroblast compartment, which exhibited the highest expression of PI16 (Supplementary Figure 2C), a marker previously associated with homeostatic fibroblast progenitor populations across organs [40]. Using this population as the putative starting state revealed a progressive fibroblast activation trajectory characterized by a transition from a quiescent resident stromal state toward an increasingly activated, matrix-remodeling phenotype. Early pseudotime cells were enriched for canonical homeostatic fibroblast markers, including PI16, DCN, and GSN, consistent with a resident fibroblast identity. Along the inferred trajectory, fibroblasts acquired a more activated transcriptional program marked by increased expression of TIMP1, FN1, SPARC, and THBS2, indicating extracellular matrix remodeling and stromal activation. This transition culminated in more differentiated fibroblast states, including COL1A1^hi^/POSTN^hi^ fibroblasts and a distinct NOTCH3^+^ fibroblast subset, suggesting divergence into specialized activated fibroblast populations. In contrast, the myofibroblast cluster formed a separate transcriptional island characterized by strong ACTA2 and TAGLN expression and was not incorporated into the main pseudotemporal path, consistent with a distinct contractile fibroblast subpopulation rather than a stepwise intermediate along the principal fibroblast differentiation trajectory (Supplementary Figure 2).

### Fb microenvironments in human acute repair and chronic fibrotic remodeling

Using BANKSY, we resolved six spatially informed immune cell subpopulations, annotated on the basis of differential marker expression and neighborhood transcriptional programs, comprising SPP1^+^ MФ, APOE^+^ MФ, C1Q^+^ MФ, monocytes, T cells, and plasma cells (Supplementary Figure 3A, C). Among the MФ subpopulations, SPP1^+^ MФ predominated in AMI with increased numbers as compared to conditions with chronic fibrotic remodeling. APOE^+^ MФ were associated with ischemic injury and were enriched in both AMI and ICM. Chronic fibrotic conditions demonstrated predominance of C1Q^+^ MФ, which were increased in chronic conditions compared to AMI (Supplementary Figure 3B, D).

We next characterized the local microenvironments of individual Fb subpopulations and immune cell populations. Clustering cells according to the composition of their immediate neighbors identified eight different multicellular spatial domains, termed cellular neighborhoods (CNs): COL1A1^hi^/POSTN^hi^/NOTCH3^+^ Fb CNs, TIMP1^hi^ Fb CNs, GSN^hi^/DCN^hi^ Fb CNs, AP1^hi^ Fb CNs, C1Q^+^ MФ CNs, APOE^+^ MФ CNs, cardiomyocyte CNs and an additional unclassified CN (Figure 4A).

**Figure 4:**
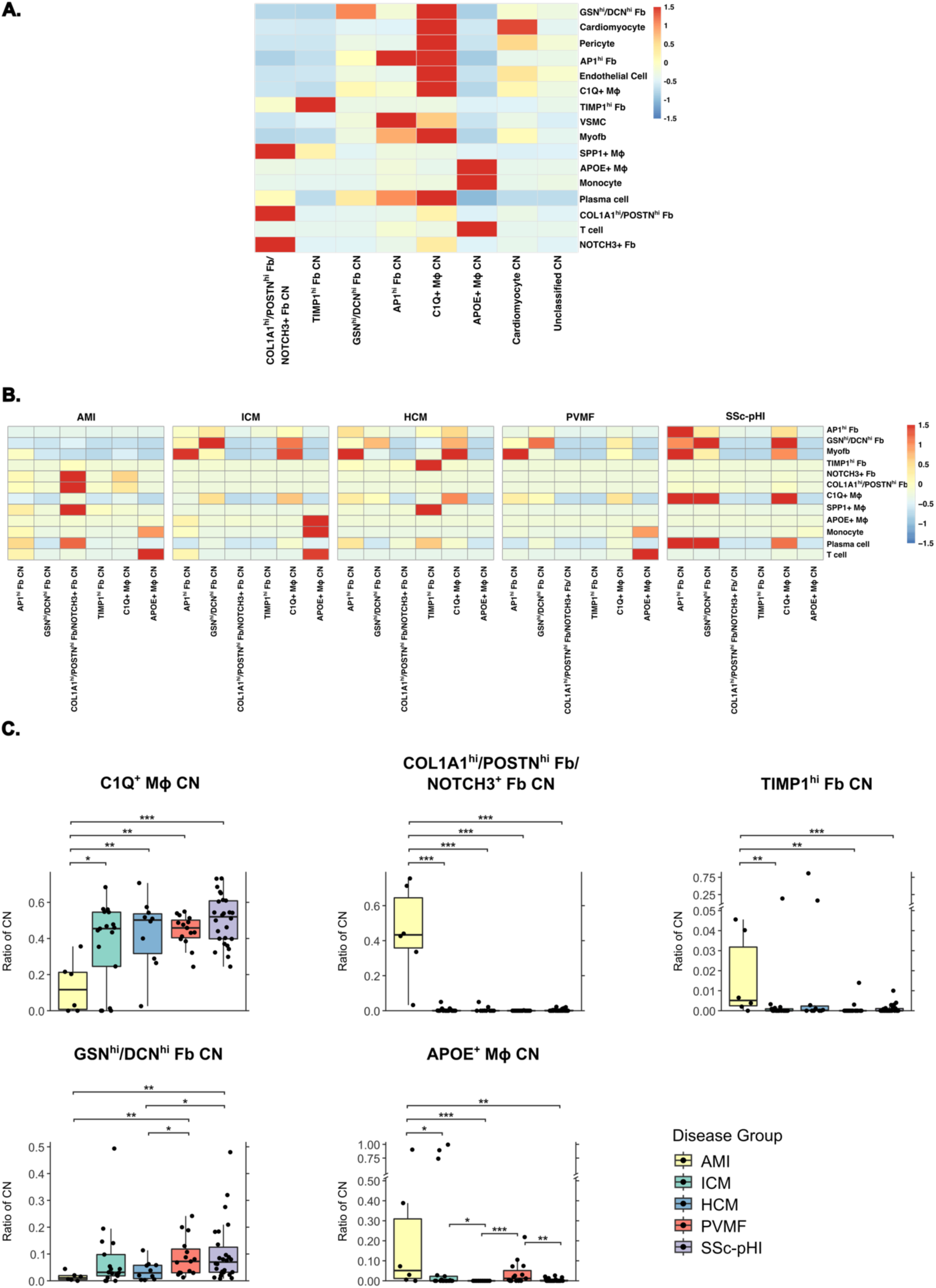
Cellular neighborhood composition and disease-associated spatial organization in myocardial biopsies. **(A)** Heatmap with the composition of the identified cellular neighborhoods. **(B)** Heatmaps showing the composition of cellular neighborhoods across disease states; values were normalized for each cell type across all cellular neighborhoods. **(C)** Box plots showing the distribution of cellular neighborhoods across disease conditions. Each dot represents an individual sample. Statistical significance (Kruskal-Wallis test) is indicated as *p < 0.05, **p < 0.01, and ***p < 0.001.

Four CNs represented fibroinflammatory niches enriched in distinct Fb and immune-cell subpopulations, namely COL1A1^hi^/POSTN^hi^/NOTCH3^+^ Fb CNs, GSN^hi^/DCN^hi^ Fb, AP1^hi^ Fb CNs, and C1Q^+^ MФ CNs. Each of these neighborhoods demonstrated specific cellular compositions of Fbs and immune cells. The COL1A1^hi^/POSTN^hi^/NOTCH3^+^ Fb CN was also enriched in SPP1^+^ MФ, which have been characterized as a key profibrotic MФ population across different fibrotic organs [41–45]. The GSN^hi^/DCN^hi^ Fb CN also demonstrated increased numbers of C1Q^+^ MФ and plasma cells. The AP1^hi^ Fb niche was co-enriched in myofibroblasts, VSMCs and plasma cells. The C1Q^+^ MФ CN included a broader range of cell types, with particular co-enrichment in AP1^hi^ Fbs, GSN^hi^/DCN^hi^ Fbs, myofibroblasts and vascular cells, but also plasma cells (Figure 4A).

The APOE^+^ MФ CN was dominated by immune cells of the monocytic lineage and T cells. In contrast, the TIMP1^hi^ Fb CN consisted almost exclusively of TIMP1^hi^ Fbs with only few other cell types. The cardiomyocyte CN also did not show enrichment of immune cells (Figure 4A).

Of note, several of these cellular neighborhoods colocalized with specific anatomical regions as defined by histology (Supplementary Figure 4B). COL1A1^hi^/POSTN^hi^/NOTCH3^+^ Fb CNs commonly localized to the infarction-associated scar. Comparison with the spatial transcriptomic reference of Kuppe et al. [3] revealed predominant correspondence of our AMI samples and cellular neighborhoods to the ischemic zone (IZ). All six samples had their highest regional score for the IZ (Supplementary Figure 5A). Mean scores across eligible samples likewise favored the IZ for all seven defined CNs, with the highest mean IZ score in the COL1A1^hi^/POSTN^hi^/NOTCH3^+^ Fb CN (Supplementary Figure 5B). This neighborhood favored the IZ in all evaluable samples. At the ROI level, 66 of 68 evaluable ROIs (97.1%) favored the IZ and only two favored the border zone (Supplementary Figure 5C).

GSN^hi^/DCN^hi^ Fb CNs localized predominantly to areas of subendocardial fibrosis and, to a lesser extent, perivascular and interstitial fibrosis. AP1^hi^ Fb CNs localized primarily to perivascular fibrosis and to a lesser extent to areas of subendocardial fibrosis (Supplementary Figure 4B).

These CNs demonstrated differences in frequencies, in cellular composition and in functional enrichment across the individual diseases (Figure 4B, C and Supplementary Figure 4A). COL1A1^hi^/POSTN^hi^/NOTCH3^+^ Fb, TIMP1^hi^ Fb, and APOE^+^ MФ CNs were enriched in acute reparative fibrosis relative to diseases with chronic fibrotic remodeling of the myocardium (Figure 4C).

Disease-specific changes in the cellular composition of these neighborhoods included enrichment of GSN^hi^/DCN^hi^ Fbs, AP1^hi^ Fbs, plasma cells, and C1Q^+^ MФ across GSN^hi^/DCN^hi^ Fbs, AP1^hi^ Fbs, and C1Q^+^ MФ fibroinflammatory niches in SSc-pHI. Another prominent disease-specific change in cellular composition included the enrichment of SPP1^+^ MФ in the TIMP1^hi^ Fb CN in HCM (Figure 4B).

Disease-associated functional differences of the individual CNs were observed in particular in AMI and SSc-pHI. Functional enrichment analysis highlighted that in AMI, the TIMP1^hi^ Fb CN was associated with ECM production and WNT signaling, whereas the same CN was preferentially linked to apoptosis in SSc-pHI. Similarly, the AP1^hi^ Fb CN was associated with collagen production in AMI, whereas it was preferentially linked to FRA signaling and T cell-mediated immunity in SSc-pHI (Supplementary Figure 4B). In the perivascular niche, pericytes were enriched for collagen assembly and biosynthesis pathways in SSc-pHI, indicating a collagen-secretory phenotype, while endothelial cells showed an immune signature through enrichment of lymphocyte mediated signaling, consistent with perivascular immune activation (Supplementary Figure 6C). Together, these findings identify CNs as disease- and context-dependent functional units of the human fibrotic myocardium.

### CODEX imaging provides orthogonal protein level support for cISH-defined myocardial cellular neighborhoods

To determine whether components of the cISH-defined cellular neighborhoods were detectable at the protein level, we analyzed 64,075 CODEX-defined cells from 31 matched donors on separate tissue sections. Manual annotation of clusters recovered the major cardiac compartments, including cardiomyocytes, ECs, mural cells, fibroblasts and macrophages (Supplementary Figure 7A, B). We further subclustered fibroblasts and macrophages and identified COL1^hi^/POSTN^hi^, GSN^hi^/DCN^hi^, AP1^hi^/TIMP1^hi^ Fbs and myofibroblasts, and C1QC^+^, SPP1^+^ and APOE^+^ macrophages, thus recapitulating major fibroblast and macrophage subpopulations described by cISH at protein level (Supplementary Figure 7 C-F).

Neighborhood analysis identified seven CNs. The COL1^hi^/POSTN^hi^ Fb CN was characterized by COL1^hi^/POSTN^hi^ Fbs and SPP1^+^ MФ. The AP1^hi^/TIMP1^hi^ Fb CN contained a prominent myofibroblast component, whereas the GSN^hi^/DCN^hi^ Fb CN also contained AP1^hi^/TIMP1^hi^ Fbs. Thus, CODEX provided orthogonal protein support for the spatial recurrence of selected neighborhood components (Figure 5).

**Figure 5:**
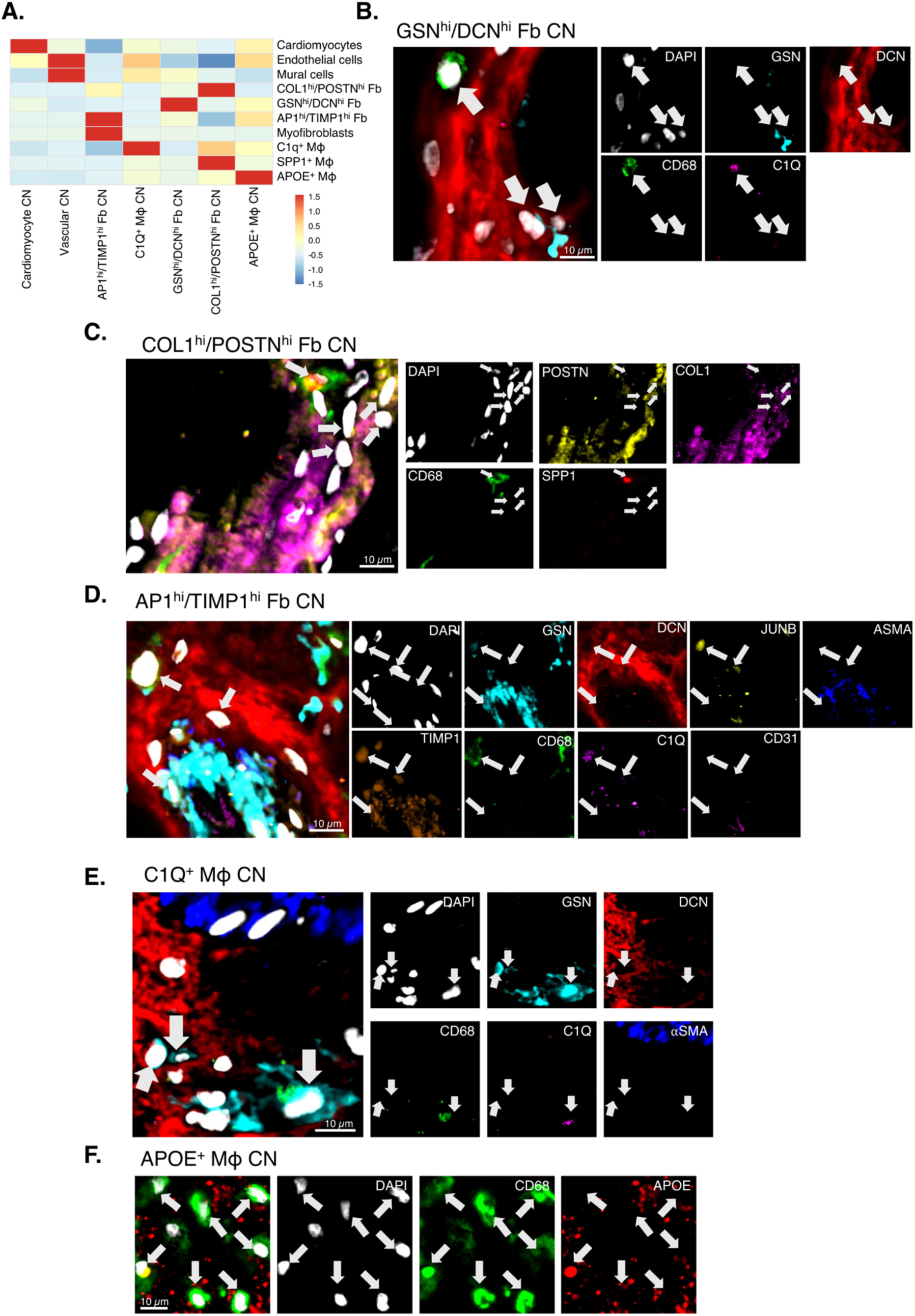
Cellular composition and representative marker distributions within fibroblast- and macrophage-enriched cellular neighborhoods in CODEX. **(A)** Heatmap showing the cell-type-scaled representation of major cardiac cell subsets and fibroblast and macrophage subpopulations across the indicated cellular neighborhoods (CNs). Red and blue denote higher and lower relative representation, respectively. **(B-F)** Representative multiplexed images and corresponding single-marker channels illustrating the cellular composition of selected CNs. **(B)** GSN^hi^/DCN^hi^ fibroblast CN, showing DAPI, GSN, DCN, CD68 and C1Q. **(C)** COL1^hi^/POSTN^hi^ fibroblast CN, showing DAPI, POSTN, COL1, NOTCH3, CD68 and SPP1. **(D)** AP1^hi^/TIMP1^hi^ fibroblast CN, showing DAPI, GSN, DCN, JUNB, TIMP1, CD68, C1Q, αSMA and CD31. **(E)** C1Q^+^ macrophage CN, showing DAPI, GSN, DCN, CD68, C1Q and αSMA. **(F)** APOE^+^ macrophage CN, showing DAPI, CD68 and APOE. Pseudocolours in the merged images correspond to those used in the respective single-marker channels. White arrows indicate representative marker-positive cells. Scale bar for reference. APOE, apolipoprotein E; CN, cellular neighborhood; COL1, collagen type I; DAPI, 4′,6-diamidino-2-phenylindole; DCN, decorin; Fb, fibroblast; GSN, gelsolin; MФ, macrophage; POSTN, periostin; SPP1, secreted phosphoprotein 1; αSMA, α-smooth muscle actin.

### Fb interaction networks in acute repair and chronic fibrotic remodeling of the myocardium

We next sought to define disease-associated changes in Fb interaction networks. Pairwise interaction analysis revealed distinct interactions in acute reparative compared with chronic fibrotic conditions. Among the interactions between Fb subpopulations, COL1A1^hi^/POSTN^hi^ and NOTCH3^+^ Fbs showed increased reciprocal interactions in AMI. By contrast, GSN^hi^/DCN^hi^ and AP1^hi^ Fbs displayed increased interactions with myofibroblasts in all chronic disease entities. Differences in Fb-to-Fb interactions were also observed within the individual chronic fibrotic diseases, in particular for SSc-pHI: GSN^hi^/DCN^hi^ Fbs demonstrated unique interactions with COL1A1^hi^/POSTN^hi^ Fbs and COL1A1^hi^/POSTN^hi^ Fbs showed increased interactions with myofibroblasts in SSc-pHI rather than with SPP1^+^ MФ, their dominant partner in AMI, indicating that the same activated Fb state operates in a different cellular context in the two conditions (Figure 6).

**Figure 6:**
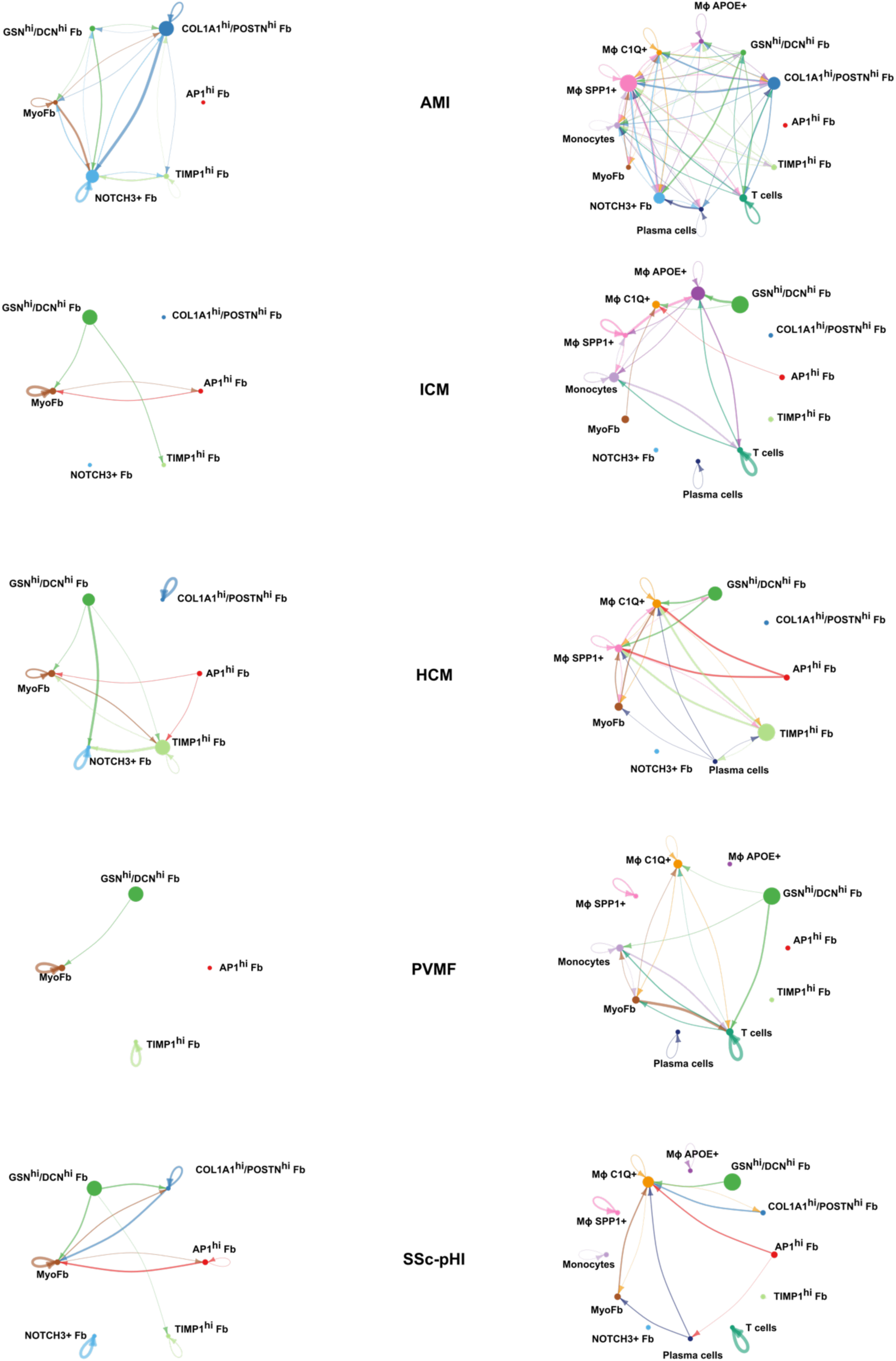
Cell-cell communication networks across disease conditions inferred by spatial ligand-receptor analysis. Circle plots depict CellChat-inferred fibroblast-fibroblast (left) and fibroblast-immune (right) interactions across AMI, ICM, HCM, PVMF, and SSc-pHI. Analyses were performed with spatial distance constraints of 400 µm for secreted signaling and 30 µm for contact-dependent signaling. Arrows indicate the direction of communication, with the arrow origin representing the sender cell population and the arrow tip representing the receiver cell population. Line width reflects the number of inferred interactions as calculated by CellChat.

Fb-immune interaction networks also differed between the different disease entities. In AMI, the most prominent Fb-immune crosstalk occurred between COL1A1^hi^/POSTN^hi^ Fbs and SPP1^+^ MФ. In chronic remodeling, Fb-immune interactions were broader and involved distinct Fb subpopulations together with additional immune cell types, including other macrophage subsets, monocytes, and T cells. Disease-specific patterns were also evident within chronic remodeling: in ICM, GSN^hi^/DCN^hi^ Fbs showed prominent interactions with APOE^+^ MФ; in HCM, both GSN^hi^/DCN^hi^ and AP1^hi^ Fbs exhibited increased interactions with C1Q^+^ and SPP1^+^ MФ; and in SSc-pHI, the strongest Fb-immune interaction was observed between GSN^hi^/DCN^hi^ Fbs and C1Q^+^ MФ. By contrast, in PVMF, GSN^hi^/DCN^hi^ Fbs interacted predominantly with monocytes and T cells rather than macrophages (Figure 6).

These disease-specific interaction patterns were accompanied by distinct ligand-receptor programs (Supplementary Figure 8). SPP1^+^ MФ in AMI and HCM, APOE^+^ MФ in ICM, and C1Q^+^ MФ in SSc-pHI showed homotypic galectin signaling and received collagen signals from COL1A1^hi^/POSTN^hi^ Fbs in AMI or GSN^hi^/DCN^hi^ Fbs in HCM, ICM, and SSc-pHI. In addition, APP signaling was consistently directed from myofibroblasts toward C1Q^+^ MФ across all chronic conditions, whereas in AMI it originated from NOTCH3^+^ Fbs toward SPP1+ MФ (Supplementary Figure 8).

Together, these data identify an infarction-associated SPP1^+^ MФ-Fb interaction axis in acute repair and more heterogeneous Fb-myeloid and Fb-lymphoid interaction circuits across chronic fibrotic remodeling.

### Spatially informed Fb subpopulations and their niches associate with clinical and histopathological features in SSc-pHI

We next assessed whether fibroblast abundance and spatial interactions were associated with clinical features in patients with SSc-pHI. In SSc-pHI patients with left ventricular dilation, COL1A1^hi^/POSTN^hi^ Fbs were more abundant than in those without left ventricular dilation (13.39% [5.81–20.20%] vs. 0.00% [0.00–1.76%], p=0.004) and showed increased interactions with cardiomyocytes (11.61% [5.81–17.37%] vs. 0.00% [0.00–0.92%], p=0.001) (Supplementary Figure 9A, B). Consistently, interactions between COL1A1^hi^/POSTN^hi^ and GSN^hi^/DCN^hi^ Fbs correlated positively with NT-proBNP levels (R=0.51, p=0.011) (Supplementary Figure 9C).

Consistent with the spatial colocalization of the GSN^hi^/DCN^hi^ Fb CN with interstitial fibrosis and its relative fibroblast-rich composition in SSc-pHI compared with other disease groups, as well as its functional enrichment for extracellular matrix-related terms, homotypic interactions among GSN^hi^/DCN^hi^ Fbs were associated with a greater histopathological extent of interstitial fibrosis (Supplementary Figure 9D). The GSN^hi^/DCN^hi^ Fb niche was also enriched in patients with anti-Scl70 autoantibodies (8.65% [6.38–18.31%] vs. 2.71% [1.37–3.38%], p=0.003), which have an increased risk of more severe cardiac involvement (Supplementary Figure 9E). Moreover, GSN^hi^/DCN^hi^ Fbs interacted more frequently with C1Q^+^ MФ in anti-Scl70-positive patients (26.51% [12.50–30.71%] vs. 9.46% [0.34–18.67%], p=0.023) (Supplementary Figure 9F). In contrast, the perivascular AP1^hi^ Fb niche was enriched in patients with digital ulcers (7.21% [4.83– 12.20%] vs. 3.23% [2.36–4.96%], p=0.008), supporting an association with the systemic vasculopathy of SSc (Supplementary Figure 9G).

## Discussion

Using spatially-resolved single-cell transcriptomics across different diseases with either repair in response to acute ischemia or chronic fibrotic remodeling due to different underlying courses, we defined a spatial taxonomy of fibroblasts, their local CNs, and fibroblast-immune cell interaction networks in human myocardial fibrosis. To our knowledge, this study is the first to compare acute repair and chronic human myocardial fibrosis across etiologies, and to provide spatially resolved single cell transcriptomic data in SSc-pHI and PVMF.

Three main conclusions emerged from our study: First, acute repair and chronic fibrosis differ widely in transcriptional pathways and cellular players, providing evidence for biologically distinct underlying remodeling programs. Second, chronic myocardial fibrosis was not uniform across etiologies but instead demonstrated different disease-associated fibroinflammatory circuits. Third, incorporation of spatial context enabled identification of specific Fb subpopulations such as AP1^hi^ and TIMP1^hi^ Fbs that were not readily separable by transcript abundance alone and lacked a discrete counterpart in reference atlases, and facilitated characterization of specific cellular interactions within local niches. These findings extend prior human cardiac atlases showing marked stromal and immune heterogeneity and are consistent with the premise that neighborhood-aware methods can uncover tissue states that approaches without spatial resolution incompletely capture [1,3,5,6]. Orthogonal CODEX protein multiplex imaging reinforced this model. Across matched donors, CODEX recovered broad cardiac compartments and fibroblast-myeloid organization. These included COL1^hi^/POSTN^hi^ fibroblast domains containing SPP1^+^ macrophages, GSN^hi^/DCN^hi^ fibroblast domains, AP1^hi^/TIMP1^hi^ fibroblast-myofibroblast domains, and C1QC^+^ or APOE^+^ macrophage domains.

In AMI, COL1A1^hi^/POSTN^hi^ and NOTCH3^+^ Fbs localized to infarction-associated CNs together with SPP1^+^ and APOE^+^ MФ. This architecture is concordant with recent findings from spatially resolved multi-omics in samples collect after myocardial infarction showing activated POSTN/COL1A1/FN1-enriched Fbs in close proximity to injury-associated myeloid populations, particularly SPP1^+^ MФ, within ischemic and border zones [3,46]. Our exploratory comparison with the regional expression signatures of Kuppe et al. [3] further supported an ischemic-zone-like transcriptional context for the sampled AMI tissue and its cellular neighborhoods. The COL1A1^hi^/POSTN^hi^/NOTCH3^+^ Fb CN showed the strongest mean ischemic-zone correspondence, while consistent correspondence across most sampled ROIs supported the broader regional context of this reparative niche. Our data extend that framework by separating an SPP1^+^ injury-remodeling MФ state from an APOE^+^ ischemia-associated MФ. Moreover, we show that these populations occupy distinct reparative niches. This supports a model in which human scar formation is orchestrated through spatially restricted Fb-MФ circuits rather than by uniform tissue-wide response.

By contrast, chronic fibrotic remodeling was dominated by GSN^hi^/DCN^hi^ Fbs, AP1^hi^ Fbs, myofibroblasts, and C1Q^+^ MФ. The relative contribution of these subpopulations differed between the individual chronic conditions. In ICM, the prominent enrichment of APOE^+^ MФ and their interaction with GSN^hi^/DCN^hi^ Fbs is consistent with persistent ischemic remodeling and immune-metabolic activation in failing myocardium [4]. In HCM, the coexistence of activated Fb subpopulations with both SPP1^+^ and C1Q^+^ MФ fits with emerging evidence that chronic leukocyte infiltration and immune activation are integral to HCM remodeling [5,6]. In PVMF, the preferential interaction of GSN^hi^/DCN^hi^ Fbs with monocytes and T cells supports a concept, in which fibrotic remodeling after viral myocarditis remains coupled to persistent immune activation even after the acute infectious phase [9,10,47]. SSc-pHI demonstrates particularly pronounced increases in GSN^hi^/DCN^hi^ Fb and C1Q^+^ MФ niches and these niches show increased numbers of plasma cells. The enrichment of plasma cells in SSc aligns with a central role of plasma cells as drivers of disease progression in SSc, as highlighted by recent reports on the clinical efficacy of plasma cell targeting approaches [48]. Collectively, these comparative data across disease entities indicate that chronic fibrotic remodeling of the myocardium shares cellular stromal responses that are adapted by disease-specific immune contexts [4–6,9,10,49].

SSc-pHI was distinguished by a higher frequency of the infarct scar-associated COL1A1^hi^/POSTN^hi^ Fb state compared with the other chronic fibrotic diseases. This TGFβ-high, matrix-producing population dominates scar formation after AMI, where it is transient and declines with scar maturation [3,46]. In SSc-pHI, COL1A1^hi^/POSTN^hi^ Fbs interacted directly with GSN^hi^/DCN^hi^ Fbs, an interaction not observed in the other chronic conditions, and were associated with distinct clinical features compared with the other fibroblasts states enriched in chronic conditions: COL1A1^hi^/POSTN^hi^ Fbs associated with cardiac dysfunction, GSN^hi^/DCN^hi^ niches with interstitial fibrosis burden and anti-Scl70 autoantibodies. This pattern indicates ongoing fibrogenesis superimposed on established fibrosis. Whether the COL1A1^hi^/POSTN^hi^ state in SSc-pHI reflects failure of resolution or continuous de novo activation sustained by recurrent microvascular injury and the plasma cell-rich immune context [48] cannot be resolved with cross-sectional data. Its association with ventricular dysfunction nevertheless nominates it as a candidate biopsy-based marker of active disease and as a therapeutic target.

Comparative analyses of histological, cellular and molecular changes demonstrated that classical histopathological patterns of fibrosis mapped onto distinct cellular microenvironments. Areas of infarction-associated scarring corresponded to COL1A1^hi^/POSTN^hi^/NOTCH3^+^ Fb CNs. In contrast, subendocardial fibrosis mapped predominantly to GSN^hi^/DCN^hi^ Fb CN, and perivascular fibrosis mapped to AP1^hi^ Fb CN. These findings highlight that myocardial remodeling is coordinated by specific fibrotic tissue niches that promote different histological patterns of fibrosis. This interpretation is aligned with recent work showing that vascular, mural, immune, and stromal cells form tightly organized fibrotic microenvironments [3].

Of interest, the frequency of selected cellular subpopulations, spatial organization and cellular interactions were associated with clinical outcomes in patients with SSc-pHI. COL1A1^hi^/POSTN^hi^ Fbs associated with cardiac dysfunction (NT-proBNP levels and left ventricular dilation) and GSN^hi^/DCN^hi^ Fb niches with anti-Scl70 autoantibodies and severity of interstitial myocardial fibrosis. The association of AP1^hi^ Fb niches with digital ulcers is consistent with the perivascular localization of these niches and may provide evidence for a vascular cell – fibroblast crosstalk that extends beyond the myocardium. Although confirmation in additional cohorts is required, these findings provide evidence that molecular and cellular patterns on myocardial biopsies may support the stratification of patients into clinically relevant phenotypes.

Our study has limitations. Normal myocardium was not included, preventing the detection of general pathophysiologic changes across disease entities. The sample size within disease groups was limited, although this is an inherent challenge of biopsy-based human myocardial studies. CODEX was performed on matched-donor tissue sections using a targeted protein panel, limiting the analysis to orthogonal support for panel-measurable neighborhood components. Moreover, cellular interaction and ligand-receptor analyses infer spatial proximity and signaling potential, respectively, rather than direct causal communication. Patients with SSc-pHI had ongoing immunosuppressive treatment at the time of biopsy, which may have also impacted the cellular composition and pathway enrichment in these samples.

Our study provides a spatial framework for human myocardial fibrosis across disease entities. By resolving context-specific fibroblast subpopulations, cellular interactions and tissue niches, these data nominate fibroblast-immune circuits for functional and translational validation and may inform the development of targeted therapeutic strategies for cardiac fibrosis.

## Acknowledgments

We gratefully acknowledge Christoph Liebel, Philipp Steinbrecher, and Lukas Sokolowski for excellent technical assistance, as well as Michael Bailey, Trieu My Van, and Michael Leon of NanoString Technologies (now Bruker Spatial Biology) for their valuable support with the CosMx runs. This work was supported by the de.NBI Cloud within the German Network for Bioinformatics Infrastructure (de.NBI) and ELIXIR-DE (Forschungszentrum Jülich and W-de.NBI-001, W-de.NBI-004, W-de.NBI-008, W-de.NBI-010, W-de.NBI-013, W-de.NBI-014, W-de.NBI-016, W-de.NBI-022).

## Disclosures

JHWD has received grants or holds contracts from: AbbVie, Agomab, Anamar, Argenx, ARXX, BMS, Bayer Pharma, Boehringer Ingelheim, Candid, Cantargia, Celgene, CSL Behring, Elli Lilly, ExoTherapeutics, GSK, Lassen, Kyverna, Mestag, Novartis, Otsuka, Sanofi-Aventis, RedX and UCB; consulting fees from: AbbVie, Anamar, ARXX, AstraZeneca, Boehringer Ingelheim, Celgene, Genentech/Roche, GSK, Novartis, Pfizer, Roche, Sulis, Spica, Tyra, UCB, Xenas; Payment or honoraria for lectures, presentations, speakers bureaus, manuscript writing or educational events from: AbbVie, Anamar, ARXX/Caluna, AstraZeneca, Boehringer Ingelheim, Elli Lilly, Genentech/Roche, GSK, Merck, Novartis, and UCB; Payment for expert testimony from: AbbVie, Anamar, ARXX/Caluna, AstraZeneca, Bayer Pharma, Boehringer Ingelheim, Cantargia, Genentech, Quell, Kyverna, Novartis, Prolium, Roche and UCB; Support for attending meetings and/or travel from: AbbVie, SOBI; Stock or stock options from: 4D Science; FibroCure. ME has received consulting fees from Ferring, AstraZeneca, Janssen (Johnson & Johnson), Genomic Health, Diaceutics, Gilead, MSD, Merck, and Bicycle Therapeutics. He has received honoraria for lectures, presentations, and educational activities from AstraZeneca, Roche, Janssen (Johnson & Johnson), Owkin, Diaceutics, Ferring, Eisai, Merck, Bicycle Therapeutics, and Zytomed Systems. He has received payments for expert testimony from AstraZeneca, Janssen (Johnson & Johnson), Astellas, Genomic Health, Diaceutics, Gilead, MSD, Merck, and Bicycle Therapeutics. He has received travel support from AstraZeneca, Roche, MSD, Janssen (Johnson & Johnson), Diaceutics, Astellas, Ferring, Eisai, Merck, and Bicycle Therapeutics. He serves on the Clinical Advisory Board of Bicycle Therapeutics and on the boards of the BRIDGE Consortium, the European Society of Uropathology (ESUP) of the European Association of Urology (EAU), and the International Society of Urological Pathology (ISUP) of the European Association of Urology (EAU). He has also received equipment, materials, or other services including research funding from STRATIFYER, Janssen (Johnson & Johnson), AstraZeneca/MedImmune, Owkin, Gilead, and Bicycle Therapeutics, BMS, QUiP GmBH. He declares stock ownership of BicycleTx and MSD Shares. KK received speaker fees from Alnylam, Pfizer and Bayer.

Claude (Anthropic) was used solely to improve the grammar and readability of the manuscript, as English is not the authors’ native language.

The other coauthors have no additional disclosures pertaining to this manuscript.

## Data and Code Availability

The data and code that support the findings of this study are available from the corresponding authors upon reasonable request.

## Non-standard Abbreviations and Acronyms

ACA: anti-centromere antibodies
ACC: American College of Cardiology
ACR: American College of Rheumatology
AMI: acute myocardial infarction
ANA: antinuclear antibodies
AP-1: activator protein 1
APP: amyloid precursor protein
ARA: anti-RNA-polymerase-III antibodies
ATA: anti-topoisomerase-I antibodies
BANKSY: Building Aggregates with a Neighborhood Kernel and Spatial Yardstick
BMI: body mass index
cISH: cyclic in situ hybridization
CK: creatine (phospho)kinase
CKD: chronic kidney disease
CMR: cardiac magnetic resonance
CN: cellular neighborhood
CODEX: co-detection by indexing
CRP: C-reactive protein
CYC: Cyclophosphamide
DAPI: 4′,6-diamidino-2-phenylindole
dcSSc: diffuse cutaneous systemic sclerosis
DLCO_SB: single-breath diffusing capacity of the lung for carbon monoxide
DLCO_VA: diffusing capacity of the lung for carbon monoxide per alveolar volume (transfer coefficient)
DNA: deoxyribonucleic acid
DU: digital ulcers
EC: endothelial cell
ECM: extracellular matrix
EDTA: ethylenediaminetetraacetic acid
eGFR: estimated glomerular filtration rate
EMB: endomyocardial biopsy
EULAR: European Alliance of Associations for Rheumatology
Fb: Fibroblast
FEV1: forced expiratory volume in 1 second
FFPE: formalin-fixed, paraffin-embedded
FVC: forced vital capacity
GAVE: gastric antral vascular ectasia
GI: Gastrointestinal
GSEA: Gene Set Enrichment Analysis
HCM: hypertrophic cardiomyopathy
H&E: hematoxylin and eosin
HFpEF: heart failure with preserved ejection fraction
HFrEF: heart failure with reduced ejection fraction
HHV6: human herpesvirus 6
HLA-DR: human leukocyte antigen–DR
hsTnI: high-sensitivity troponin I
hsTnT: high-sensitivity troponin T
ICM: ischemic cardiomyopathy
IL: Interleukin
ILD: interstitial lung disease
IQR: interquartile range
kNN: k-nearest neighbors
LA: left atrium
lcSSc: limited cutaneous systemic sclerosis
LGE: late gadolinium enhancement
LV: left ventricle / left ventricular
LVEF: left ventricular ejection fraction
Mf: Macrophage
MHC: major histocompatibility complex
MMF: mycophenolate mofetil
MRI: magnetic resonance imaging
mRNA: messenger ribonucleic acid
mRSS: modified Rodnan skin score
MSK: Musculoskeletal
MTX: Methotrexate
NA: not available
NSAID: nonsteroidal anti-inflammatory drug
NT-proBNP: N-terminal pro-B-type natriuretic peptide
PAH: pulmonary arterial hypertension
PAPsys: systolic pulmonary arterial pressure
PBS: phosphate-buffered saline
PC: principal component
PCA: principal component analysis
PCR: polymerase chain reaction
PDGF: platelet-derived growth factor
PPI: proton pump inhibitor
PROGENy: Pathway RespOnsive GENes
PVMF: postviral myocardial fibrosis
QPTIFF: quantitative pathology tagged image file format
RA: right atrium
RAAS: renin-angiotensin-aldosterone system
RNA: ribonucleic acid
RT-PCR: reverse transcription polymerase chain reaction
RV: right ventricle / right ventricular
scANVI: single-cell ANnotation using Variational Inference
sc/snRNASeq: single-cell / single-nucleus RNA sequencing
SIP: Spatial Informatics Platform
SM-actin, αSMA: (α-)smooth muscle actin
SMI: spatial molecular imaging
SSc: systemic sclerosis
SSc-pHI: systemic sclerosis-associated primary heart involvement
SV: Supraventricular
TAPSE: tricuspid annular plane systolic excursion
TFR: tendon friction rubs
TGF-β: transforming growth factor-β
Tris: tris(hydroxymethyl)aminomethane
TTE: transthoracic echocardiography
UMAP: uniform manifold approximation and projection
V: ventricular
VSMC: vascular smooth muscle cell

