## Supplemental Material for "Spatial organization of fibroblast subpopulations and niches across acute repair and chronic fibrotic remodeling of the human myocardium"

### Supplemental Publication Material

#### Supplemental Methods

##### Patients and endomyocardial samples

Demographic and clinical characteristics of SSc patients are presented in Supplementary Table 1. Demographic and clinical characteristics of PVMF, ICM, HCM, and AMI patients are presented in Supplementary Table 2. All subjects were of European ancestry. This study was conducted in accordance with the Declaration of Helsinki. Retrospective data collection and anonymized analysis of the patients were conducted in accordance with local government law without the requirement for informed consent. The study was approved by the ethical committees of the Heinrich Heine University Düsseldorf (Study Number 2023-2561), the Friedrich Alexander University Erlangen (Approval Nr. 4607) and the Institutional Review Board of the Eberhard Karls University Tübingen (411/2021BO2).

All SSc patients fulfilled the American College of Rheumatology (ACR)/ European Alliance of Associations for Rheumatology (EULAR) 2013 classification criteria for SSc [15]. Early SSc was defined as a disease duration of less than three years from the first non-Raynaud disease manifestation. SSc patients were classified as inflammatory if one or more of the following parameters were present at the time of biopsy: CRP > 0.6 mg/dl, thrombocyte count over 330,000/µl, arthritis (clinically apparent or synovitis on joint ultrasound or MRI examination), or myositis (increased CK associated with clinical signs of myositis and/or electromyographic or MRI features). The presence of heart failure and its subtypes were assessed using the American College for Cardiology (ACC) criteria [16]. The left ventricular ejection fraction (LVEF) was measured either by transthoracic echocardiography (TTE) or cardiac magnetic resonance (CMR).

Patients were classified as PVMF when endomyocardial biopsy had been performed for clinically suspected chronic myocarditis - unexplained cardiac symptoms, newly documented arrhythmia, or echocardiographic abnormality without an alternative cardiac diagnosis - and the biopsy showed interstitial myocardial fibrosis in a diffuse and/or focal, in part perivascular distribution, without the confluent subendocardial or transmural scarring typical of ischemic injury. Classification additionally required either detection of a cardiotropic viral genome in myocardial tissue by nested (RT-)PCR or a histological and immunohistological constellation compatible with healed or chronic postviral myocarditis; biopsies showing giant-cell or eosinophilic myocarditis, cardiac amyloidosis, sarcoidosis or storage disease were not assigned to this group. The PCR-negative samples were classified on the basis of the histopathological pattern. Because cardiotropic viral genomes may persist in the myocardium without ongoing virus-mediated injury, and a negative PCR does not exclude a previously cleared infection, viral genome detection was regarded as supportive rather than defining and PVMF is therefore used as a clinicopathological designation. Group assignment was made by an experienced cardiopathologist together with the treating cardiologist.

ICM without myocardial infarction was defined as left ventricular systolic dysfunction in the presence of coronary artery disease of sufficient extent to account for the ventricular dysfunction, in the absence of a documented previous or acute myocardial infarction [17].

HCM was defined according to the 2023 European Society of Cardiology Guidelines for the management of cardiomyopathies as a maximal left ventricular wall thickness ≥15 mm in any myocardial segment that was not explained solely by abnormal loading conditions [18].

AMI was defined according to the Fourth Universal Definition of Myocardial Infarction, as adopted by the 2023 ESC Guidelines for the management of acute coronary syndromes, by a rise and/or fall in cardiac troponin with at least one value above the 99th-percentile upper reference limit together with clinical evidence of acute myocardial ischemia. Myocardial tissue was obtained within 14 days of the index infarction [19,20].

**Sample processing**

Ten µm thick FFPE sections were subjected to imaging-based spatial transcriptomic analysis to obtain single-cell transcriptional data while preserving the spatial context of the cells within each biopsy [21]. cISH was conducted in Seattle, WA, USA, by NanoString Technologies (now Bruker Spatial Biology Inc., Seattle, WA, USA). The analysis utilized a pre-existing mRNA probe panel comprising 950 genes (The Human Universal Cell Characterization RNA Core Panel), supplemented with 19 additional probes against genes characteristic for specific fibroblast subpopulations. The final probe panel (Supplementary Table 3) also included ten negative control probes targeting sequences from the External RNA Controls Consortium [22]. Cell segmentation was performed on the CosMx SMI platform, employing an automated pretrained model based on the Cellpose algorithm, which utilizes cellular membrane, nuclear, and RNA stainings. Transcript localization data and single-cell expression matrices were extracted from the AtoMx SIP platform using the Export and Flat File Export modules.

**Quality control and data normalization**

The files from AtoMx were imported as a SpatialFeatureExperiment (.sfe file) object using the Voyager R package. Poorly segmented objects and artifacts were excluded based on DAPI staining intensity and object size using predefined criteria. Cells with low RNA counts (≤6) and a high proportion of negative control counts (≥10%) were excluded from subsequent analyses.

Expression values were normalized by dividing each cell's raw RNA counts by the total RNA counts per cell, applying a lower threshold of 20 to prevent excessive upscaling of low-count cells. A z-score-scaled count matrix was then generated by performing gene-wise normalization on the normalized expression data.

**Spatially informed phenotyping**

Identification of cellular populations using both gene expression profile and spatial information was performed via spatial clustering using the Banksy R package [23]. A Building Aggregates with a Neighborhood Kernel and Spatial Yardstick (BANKSY) expression matrix was constructed. This matrix type not only incorporated each cell's individual expression profile, but also the expression profiles of its cellular neighbors before performing dimensionality reduction. Each individual cell's neighbor expression profiles were constructed using two parameters: 1) The weighted mean expression based on the distances between cells. The weight decreases exponentially with distance and is normalized by the median distance among the neighbors. 2) The gradient expression, which is similar to the mean expression but additionally includes an azimuthal component which accounts for the directional information from the neighbors. The neighboring cells were defined by k-nearest neighbors (kNN) with k=15 for the mean expression and k=30 for the gradient expression.

A neighbor-augmented expression was generated by combining the cells' own expression and two neighbor expression matrices with the spatial mixing factor $\lambda = 0.2$. Non-spatial clustering was also performed by setting $\lambda = 0$ for comparison. To identify cell populations, we further performed dimensionality reduction by PCA and Leiden clustering with the first 20 PC. Uniform Manifold Approximation and Projection plots (UMAPs) were created from the first 20 PC for the data visualization.

**Pseudotime trajectory inference**

Pseudotime trajectories describing fibroblast differentiation were generated using the R package Monocle (v3) [24,25]. For trajectory inference, the same assay and reduced dimensional embeddings used for BANKSY clustering were applied. Cells were ordered along pseudotime using the order_cells function, with $GSN^{hi}/DCN^{hi}$ Fb designated as the root population.

**Cellular neighborhoods and interaction analysis**

Cellular neighbors were defined as adjacent cells identified through Delaunay triangulation, with centroid distances within 48 μm. Neighbor cell pairs were detected using imcRtools R package [26]. To investigate the cellular local environments, we identified CNs using the previously described protocols [22,27,28]. We first computed the composition of neighboring cells for each cell, and K-means clustering was applied to these compositions to define CNs. The cells residing in the same CN shared similar neighboring cell types. We created heatmaps to visualize the cell composition within CNs. The heatmaps were scaled according to the proportion of individual cell type in the entire dataset, unless otherwise specified.

To analyze the spatial proximity between cells, we performed cellular pairwise interaction analysis using the imcRtools R package [26]. The significance of individual pairwise interactions was determined using permutation testing. These tests calculated the number of neighboring cells for each subpopulation and compared it to an empirical null distribution obtained by randomly shuffling cell type labels for 1000 permutations. A pairwise interaction with a p-value below 0.05 was considered as significant. To account for the heterogeneity among donors, the permutation testing was performed within each sample independently, i.e. the annotations were shuffled only among cells from the same individual. We interpreted statistically significant higher observed cell counts as cellular attraction and lower counts as cellular avoidance. Dot plots and interaction network visualizations were generated using ggplot2 and ggraph R packages.

**Functional analysis**

Normalized count data was used for expression profiling. Marker gene expression for each cell population was visualized using heatmaps and UMAP plots. The marker genes were discovered by Wilcoxon rank sum tests implemented in the Seurat R package with the FindAllMarkers function. Smoothed expression values were calculated using k-NN in the UMAP space, following the instruction from Bruker Spatial Biology (https://nanostring-biostats.github.io/CosMx-Analysis-Scratch-Space/) for the visualization of gene expression on UMAP plots. To characterize fibroblast populations, we computed expression scores for extracellular matrix components, including collagen, using the AddModuleScore function in the Seurat R package. The ECM geneset used for the scores was obtained from the Core Matrisome Annotations. To explore the biological functions across populations and CNs, we generated pseudo-bulk expression profiles by averaging expression within defined cell populations or conditions. The pathway activity scores were computed by fgsea using decoupleR R package, with a curated gene set obtained from the Bader Lab, available in the GSEA database of the Broad Institute, Inc., Massachusetts Institute of Technology and Regents of the University of California (version Human_GOBP_AllPathways_noPFOCR_no_GO_iea_March_01_2025). TGF-β activity score was inferred using a multivariate linear model with PROGENy gene signatures. The top 2000 gene weights were retrieved from PROGENy using the OmnipathR R package. The gene sets and weights were filtered to match the genes included in the CosMx panel for the functional analysis. To investigate cell-cell communication, we conducted ligand-receptor analysis using the CellChat R Package. Ligand-receptor interaction data was sourced from the CellChatDB database. The averaged expression was computed by truncated mean, with trimming proportion set to 10%. We set distance constraints of 400 μm for secreted signaling and 30 μm for cell contact-dependent signaling. Interactions involving fewer than five cells were excluded from further analysis to ensure robust communication detection.

**Integration of cISH data with publicly available sc/snRNASeq data**

To enable comparison with previously described fibroblast populations, our dataset was integrated with publicly available single-cell/single-nucleus RNA sequencing (sc/snRNASeq) datasets [1,3,29,30]. Raw count matrices and accompanying metadata were obtained from the original repositories or supplementary materials. When cell-type annotations were available, these were retained as provided. In cases where nomenclature differed between studies, labels were harmonized based on the marker genes and cell-type definitions reported in the respective publications to ensure consistency across datasets. All analyses were performed in Python using the scvi-tools framework. Prior to integration, genes were matched across datasets based on shared gene identifiers, and count matrices were restricted to the common gene set. Quality-control filtering and preprocessing were performed as described in the preprocessing section above. Integration and label transfer were performed using single-cell ANnotation using Variational Inference (scANVI). Annotated cells from the reference datasets were used to train the model, while cells from our dataset were treated as unlabeled query cells. The model jointly learns a shared latent representation of all cells and a classifier that predicts cell-type labels for unlabeled cells, providing both predicted labels and associated prediction probabilities. The learned latent space was used for downstream analyses and visualization. To assess integration quality and annotation consistency, several complementary visualizations were generated. Two-dimensional embeddings of the scANVI latent space were computed using UMAPs and colored by dataset of origin, predicted cell type, or clustering results to evaluate mixing of cells across datasets and separation of biologically meaningful populations. Prediction probabilities produced by scANVI were examined to assess the confidence of label transfer and to identify populations with ambiguous or transitional identities. Sankey diagrams were generated to visualize the correspondence between cell populations identified in this study and Fb subtypes predicted by scANVI. For each cell, the cluster or annotation derived from our analysis was paired with its predicted label. These counts were used to define the links in the Sankey diagrams, where the width of each connection is proportional to the number of cells shared between populations, enabling visual assessment of how clusters map to reference-defined cell types (Supplementary Figure 4).

**Transcriptomic correspondence to myocardial infarction regions**

To assess which infarct region our AMI samples most closely resembled, we compared their transcriptomic profiles with the remote-zone (RZ), border-zone (BZ), and ischemic-zone (IZ) reference data from Kuppe et al. [3] (Zenodo record 6578047). Raw counts from 17 Visium libraries were summed by reference patient and region, yielding 15 profiles from 10 patients; control and fibrotic-zone profiles were excluded. The six AMI samples were analyzed separately, with raw counts from quality-filtered cells aggregated for each whole sample, CN within each sample, and region of interest (ROI). Existing CN annotations were retained. The comparison used 666 genes shared between the cISH panel and all reference libraries. Reference regional signatures were derived using expression filtering, TMM normalization, voom weighting, and limma models with patient blocking and robust empirical Bayes moderation. Each region was compared with the equally weighted mean of the other two. Up to 25 upregulated and 25 downregulated genes per region were selected by absolute moderated t statistic, requiring $|\log 2 \text{ fold change}| \geq 0.25$ without a significance threshold. For each query aggregate, regional scores were calculated as the mean within-profile percentile rank of upregulated genes minus that of downregulated genes and standardized against reference profiles with equal weighting of the three regions. Eligible aggregates required ≥20 cells, ≥1,500 shared-gene counts, and ≥100 detected shared genes. Whole-sample scores were displayed individually, whereas CN scores were averaged with equal weight for each eligible sample. ROI correspondence was defined by the highest regional score and mapped to the mean native coordinates of retained cells within each ROI. Reference performance was assessed by leave-one-patient-out cross-validation, repeating preprocessing, model fitting, and signature selection within each training fold.

#### Co-detection by indexing (CODEX) multiplex protein tissue imaging

##### *Antibody conjugation, validation and titration*

Carrier-free antibodies were conjugated to unique DNA oligonucleotides according to an established CODEX/PhenoCycler protocol [31]. Briefly, 35-50 μg antibody was loaded onto a filter column and reduced for 30 min at room temperature in 360 μL PBS containing 2.5 mM tris(2-carboxyethyl)phosphine and 2.5 mM EDTA. The reduction reagent was removed by two washes with Buffer C containing Tris, 0.15 M NaCl, 1 mM EDTA and 0.02% sodium azide. The corresponding DNA oligonucleotide was dissolved in 400 μL PBS, added to the reduced antibody and incubated for 2 h at room temperature. Unbound oligonucleotide was removed using high-salt PBS containing 0.9 M NaCl and 0.02% sodium azide. The antibody-oligonucleotide conjugate was recovered in CODEX antibody stabilizer at 2 μL/μg antibody. Conjugated antibodies were validated and titrated before assembly of the complete panel (Supplementary Table 4).

##### *Myocardial tissue preparation and staining*

In total, 36 myocardial samples from 36 patients were analyzed by CODEX. The cohort comprised three patients with AMI, nine patients with ICM, five patients with HCM, nine patients with PVMF, and ten patients with SSc-pHI. Ten-µm FFPE myocardial tissue sections were deparaffinized in xylene and rehydrated through a graded ethanol series. Antigen retrieval was performed for 20 min in Tris–EDTA buffer, pH 9.0, using a pressure cooker. Tissue autofluorescence was reduced by placing the slides in bleaching solution containing 25 mL PBS, 4.5 mL 30% hydrogen peroxide and 0.8 mL 1 M NaOH between two high-power LED panels for 45 min; this bleaching cycle was performed twice. The sections were washed in hydration buffer containing 10 mM Tris, pH 7.5, 0.02% sodium azide, 0.1% Triton X-100, 10 mM $MgCl_2 \cdot 6H_2O$ and 150 mM NaCl. Sections were incubated in Akoya Staining Buffer for 30 min and subsequently in Akoya Blocking Buffer for 30 min. The complete DNA-conjugated primary-antibody cocktail (Supplementary Table 4) was prepared in Staining Buffer, applied to the sections and incubated overnight at 4°C on an orbital shaker. Slides were then washed and fixed according to the manufacturer's instructions and the established CODEX protocol [31,32].

***Multicycle imaging***

Multicycle imaging was performed using a PhenoCycler-Fusion 1.0 system equipped with a 20× air objective (Akoya Biosciences). Barcoded reporter oligonucleotides labelled with ATTO 550 or ATTO 647 were used to visualize up to two antibody targets per cycle, and nuclei were stained with DAPI in every cycle. The instrument generated cycle-aligned multiplex images in QPTIFF format. Images were trimmed in QuPath version 0.4.3 before single-cell feature extraction.

***Single-cell feature extraction, transformation and quality control***

The myocardial QPTIFF image was analyzed directly in QuPath. Nuclei were detected with StarDist using `StarDistSegmentation.groovy` and the pretrained single-channel fluorescence model `dsb2018_heavy_augment.pb` (QuPath [33]; StarDist [34]). QuPath-exported cell and region-of-interest tables and per-cell polygon outlines were used to construct single-cell objects. The exported measurements included detection probability, nuclear and cell morphology, cell-centroid coordinates and mean marker intensities for the nuclear, cytoplasmic, membrane and whole-cell compartments. Whole-cell mean intensities were used for downstream annotation.

The matched cross-modal analysis included 64,075 quality-controlled CODEX cells from 31 donors with corresponding cISH data. Cells were retained when their segmented area was 10-250 $\mu m^2$ and their mean nuclear DAPI intensity was at least 900. For each marker, raw whole-cell mean intensities were winsorized at the 0.1st and 99.9th percentiles calculated among quality-control-passing cells and then transformed using the inverse hyperbolic sine function with a fixed cofactor of 100. DAPI was used for nuclear segmentation and quality control but was excluded from protein-based cell annotation and spatial-program scoring.

***Clustering, cell annotation and cellular neighborhood analyses of CODEX data***

For clustering and dimensionality reduction, the arcsinh transformed value for each marker was centered on the donor-specific median and divided by the donor-specific interquartile range (R quantile type 8), then clipped to −5 to 5. DAPI was excluded. RPhenograph clustering (Rphenograph package) and UMAP (uwot package) were performed directly on the donor-scaled protein matrix without PCA.

For broad cell-family annotation, all cells were jointly clustered with RPhenograph using k = 150 and nine proteins: cardiac troponin T (TNNT2), gelsolin (GSN), decorin (DCN), NOTCH3, αSMA, CD45, CD68, CD31 and vWF. The resulting 15 communities were manually mapped, using donor-balanced marker profiles, to cardiomyocytes, endothelial cells, mural cells, fibroblasts and macrophages.

Fibroblasts were subsequently subclustered with k = 40 using GSN, DCN, periostin (POSTN), collagen I (COL1), JUNB, TIMP1, NOTCH3 and αSMA. Macrophages were subclustered with k = 200 using C1QC, osteopontin (SPP1) and APOE.

Cellular neighbors were identified with a similar approach for CODEX as for cISH (see above).

**Histopathological and immunohistological analysis**

Samples were immediately fixed under sterile conditions in 4% buffered formaldehyde for routine light microscopic evaluation of histological and immunohistological features. Paraffin-embedded tissue sections, 4 µm in thickness, were prepared as serial sections and examined by light microscopy after staining with hematoxylin and eosin (H&E), Masson's trichrome, Giemsa, and Congo red, as well as immunohistochemical staining for smooth muscle actin (SM-actin). Additional specimens were preserved in RNAlater (Ambion Inc., Foster City, CA) for RT-PCR-based detection of viral genomes. Immunohistochemistry was performed using an avidin-biotin immunoperoxidase technique according to the manufacturer's instructions (Vectastain Elite ABC Kit, Vector, Burlingame, CA). Monoclonal antibodies against CD3, CD68, and HLA-DR-α (Novocastra Laboratories, Newcastle upon Tyne, UK; DAKO, Glostrup, Denmark; DAKO, Hamburg, Germany, respectively) were used to identify and characterize mononuclear inflammatory infiltrates, including T lymphocytes, macrophages, and MHC class II expression in antigen-presenting cells. Inflammation was assessed according to the World Health

Organization/International Society and Federation of Cardiology Task Force criteria for the definition and classification of cardiomyopathies [35]: endomyocardial biopsies were considered inflamed when focal or diffuse mononuclear infiltrates of at least 14 immune cells/mm² (CD3-positive T cells and/or CD68-positive macrophages) were present in the myocardium, together with increased MHC class II expression [35]. For inflammatory heart disease, immunohistochemical subtyping of inflammatory cells was additionally performed to distinguish, for example, virus-associated myocarditis from giant cell or eosinophilic myocarditis, in line with recommendations from the Association for European Cardiovascular Pathology [36]. In this manner, standard histological and immunohistological staining methods were combined to establish the specific underlying cardiac diagnosis. The degree of inflammation was further graded using a modified scheme: grade 0, no inflammation; grade 1, isolated inflammatory cells (T lymphocytes and macrophages ≥14/mm²); grade 2, a few inflammatory foci; grade 3, multiple inflammatory foci; and grade 4, marked inflammation. Cardiac fibrosis was assessed by an experienced cardiopathologist and categorized into tertiles based on extent of fibrosis: grade 1 (0–10%), grade 2 (11–15%), and grade 3 (16–32%) [12]. Viral genome analysis was performed on RNAlater-preserved endomyocardial biopsy specimens using nested (RT-)PCR assays. The panel included enteroviruses (including coxsackieviruses and echoviruses), adenoviruses, influenza A and B viruses, parvovirus B19, human herpesviruses 6, 7, and 8, herpes simplex viruses 1 and 2, human cytomegalovirus, varicella-zoster virus, and Epstein–Barr virus. Amplification procedures were carried out in accordance with the manufacturer's instructions (AGS, Heidelberg, Germany) and established protocols described previously [12,37]. Samples were classified as virus-positive when viral nucleic acids were detected by nested (RT-)PCR and the findings were subsequently verified by sequence analysis.

#### Statistical analysis

Continuous variables were summarized as mean ± standard deviation or median and interquartile range, depending on data distribution. Normality of continuous variables was assessed using the Shapiro-Wilk test. Frequencies were reported as absolute counts and relative percentages within the cohort. Inferential statistical analyses were conducted using R [38] in RStudio. Differences in cluster frequencies (per-patient proportions) between disease conditions were evaluated using the Kruskal-Wallis test. Graphs were generated using the ggplot2 R package. Statistical significance was defined as $p < 0.05$.

#### Supplementary Tables

**Supplementary Table 1. Demographic and clinical data of the SSc cohort.**

| Parameters | Value |
|---|---|
| **Demographics** | |
| Age at biopsy, years (median, IQR) | 53 (13) |
| Male (n, %) | 6 (46.15 %) |
| Smoker (n, %) | 3 (23.07 %) |
| Ex-Smoker (n, %) | 4 (30.76 %) |
| BMI, kg/m$^2$ (median, IQR) | 23.4 (5.87) |
| Death up to Jan. 2026 (n, %) | 1 (7.69 %) |
| **Disease Characteristics** | |
| Age at First non-Raynaud Manifestation, years (median, IQR) | 48 (17) |
| Age at SSc Diagnosis, years (median, IQR) | 48 (16.25) |
| Age at SSc-pHI Diagnosis, years (median, IQR) | 54.5 (14) |
| ACR/EULAR Criteria fulfilled (n, %) | 13 (100 %) |
| Skin Involvement | |
| dcSSc (n, %) | 9 (69.23 %) |
| lcSSc (n, %) | 4 (30.76 %) |
| Progressive Skin Disease (n, %) | 6 (46.15 %) |
| mRSS at EMB (median, IQR) | 20.5 (12.25) |

| | |
|---|---|
| Peak mRSS up to EMB (median, IQR) | 21 (16) |
| MSK Involvement | |
| TFR (n, %) | 0 |
| Arthritis (n, %) | 0 |
| Myositis (n, %) | 5 (38.46 %) |
| Pulmonary Involvement | |
| ILD (n, %) | 5 (38.46 %) |
| Progressive ILD (n, %) | 1 (7.69 %) |
| FVC, % predicted (median, IQR) | 94.9 (18.15) |
| FEV1, % predicted (median, IQR) | 95.4 (10.22) |
| DLCO_SB, % predicted (median, IQR) | 61 (17.87) |
| DLCO_VA, % predicted (median, IQR) | 82.9 (21.55) |
| PAH (n, %) | 2 (15.38 %) |
| GI involvement | |
| Upper GI Tract (n, %) | 3 (23.07 %) |
| Lower GI Tract (n, %) | 1 (7.69 %) |
| Both (n, %) | 2 (15.38 %) |
| Vasculopathy | |
| Recurring Digital Ulcers (n, %) | 4 (30.76 %) |
| Telangiectasias (n, %) | 3 (23.07 %) |
| GAVE Syndrome (n, %) | 0 |
| Scleroderma Renal Crisis (n, %) | 0 |
| Comorbidities | |
| Hypothyroidism (n, %) | 3 (23.07%) |
| Diabetes mellitus (n, %) | 0 |
| Arterial Hypertension (n, %) | 1 (7.69 %) |
| Chronic Coronary Disease (n, %) | 4 (30.76 %) |
| Dyslipidemia (n, %) | 2 (15.38 %) |
| CKD (n, %) | 1 (7.69 %) |
| **Laboratory Parameters** | |
| ANA (n, %) | 11 (84.61 %) |
| ATA (n, %) | 8 (61.53 %) |
| ACA (n, %) | 0 |
| ARA (n, %) | 0 |
| Other (n, %) | 3 (23.07 %) |
| CK, U/l (median, IQR) | 244 (462) |
| hsTnI, ng/l (median, IQR) | 285 (185) |
| hsTnT, ng/l (median, IQR) | 217 (826) |
| NT-proBNP, pg/ml (median, IQR) | 726 (1,341.5) |
| eGFR, ml/min/1,73 $m^2$ (median, IQR) | 90 (1) |
| **Primary Heart Involvement** | |
| Previously Diagnosed SSc-pHI through EMB (n, %) | 0 |
| Previously Diagnosed SSc-pHI through cardiac-MRI | |
| Negative (n, %) | 7 (53.84 %) |
| Positive (n, %) | 6 (46.15 %) |
| MRI Characteristics | |
| Dilated Cavities (n, %) | 2 (15.38 %) |
| Pericardial Effusion (n, %) | 2 (15.38 %) |
| LGE (n, %) | 4 (30.76 %) |
| Diffuse Fibrosis (n, %) | 3 (23.07 %) |
| Hypokinesia (n, %) | 2 (15.38 %) |
| Indication for EMB | |
| Clinical Progression (n, %) | 3 (23.07 %) |
| Persistent Troponinemia (n, %) | 7 (53.84 %) |
| Cardiac Decompensation (n, %) | 2 (15.38 %) |
| Ventricular Arrhythmia (n, %) | 1 (7.69 %) |

| | |
|---|---|
| Ischemic Event (n, %) | 2 (15.38 %) |
| Histologic features of the EMB | |
| Chronic Myocarditis (n, %) | 13 (100 %) |
| Macrophage Dominated (n, %) | 1 (7.69 %) |
| Lymphocyte Dominated (n, %) | 12 (92.3 %) |
| Interstitial Fibrosis (n, %) | 12 (92.3 %) |
| Grade I (mild) (n, %) | 2 (15.38 %) |
| Grade II (moderate) (n, %) | 7 (53.84 %) |
| Grade III (severe) (n, %) | 3 (23.07 %) |
| Microangiopathy (n, %) | 1 (7.69 %) |
| Heart Failure | |
| HFpEF (n, %) | 5 (38.46%) |
| HFrEF (n, %) | 1 (7.69 %) |
| Arrhythmias (n, %) | 3 (23.07 %) |
| Only SV (n, %) | 0 |
| Only V (n, %) | 1 (7.69 %) |
| Both SV and V (n, %) | 2 (15.38 %) |
| Conduction Blocks (n, %) | 0 |
| Transthoracic Echocardiography Parameters | |
| LVEF, % (median, IQR) | 59 (8) |
| TAPSE, mm (median, IQR) | 28 (5.5) |
| PAPsys, mmHg (median, IQR) | 32 (28.75) |
| Diastolic Dysfunction (n, %) | 3 (23.07 %) |
| Grade 1 (n, %) | 0 |
| Grade 2 (n, %) | 2 (15.38 %) |
| Grade 3 (n, %) | 0 |
| Restrictive Dysfunction (n, %) | 0 |
| RV Dilation (n, %) | 2 (15.38 %) |
| LV Dilation (n, %) | 2 (15.38 %) |
| RA Dilation (n, %) | 0 |
| LA Dilation (n, %) | 1 (7.69 %) |
| **Treatment** | |
| Non-immunosuppressive Treatment | |
| PPI (n, %) | 10 (76.92 %) |
| Prokinetic (n, %) | 2 (15.38 %) |
| Diuretic (n, %) | 2 (15.38 %) |
| NSAID (n, %) | 3 (23.07 %) |
| Betablocker (n, %) | 2 (15.38 %) |
| Statin (n, %) | 2 (15.38 %) |
| RAAS Inhibitor (n, %) | 3 (23.07 %) |
| Vitamin D (n, %) | 4 (30.76 %) |
| Levothyroxine (n, %) | 3 (23.07 %) |
| Other Antiarrhythmic Drugs (n, %) | 1 (7.69 %) |
| PAH Therapy (n, %) | 2 (15.38 %) |
| Immunosuppressive Therapy at EMB | |
| Prednisolone (n, %) | 5 (38.46 %) |
| MMF (n, %) | 3 (23.07 %) |
| MTX (n, %) | 1 (7.69 %) |
| CYC (n, %) | 1 (7.69 %) |
| History of immunosuppressive Therapy up to EMB | |
| Prednisolone (n, %) | 6 (46.15 %) |
| MMF (n, %) | 3 (23.07 %) |
| MTX (n, %) | 1 (7.69 %) |
| CYC (n, %) | 1 (7.69 %) |
| Pacemaker or Defibrillator (n, %) | 1 (7.69 %) |

BMI – Body Mass Index, SSc – Systemic Sclerosis, SSc-pHI – Systemic Sclerosis Primary Heart Involvement, dcSSc – Diffuse Cutaneous Systemic Sclerosis, lcSSc – Limited Cutaneous Systemic Sclerosis, mRSS – Modified Rodnan Skin Score, EMB – Endomyocardial Biopsy, MSK – Musculoskeletal, TFR – Tendon Friction Rubs, ILD – Interstitial Lung Disease, FVC – Forced Vital Capacity, FEV1 – Forced Expiratory Volume in 1 second, DLCO_SB – Single-Breath Diffusing Capacity for Carbon Monoxide, DLCO_VA – Transfer Coefficient, PAH – Pulmonary Arterial Hypertension, GI – Gastrointestinal, GAVE – Gastric Antral Vascular Ectasia, CKD – Chronic Kidney Disease, ANA – Antinuclear Antibodies, ATA – Anti-Topoisomerase-I Antibodies, ACA – Anti-Centromere Antibodies, ARA – Anti-RNA-Polymerase-III Antibodies, CK – Creatine Phosphokinase, hsTnI – High Sensitivity Troponin I, hsTnT – High Sensitivity Troponin T, NT-proBNP – N-Terminal-pro-Brain Natriuretic Peptide, eGFR – Estimated Glomerular Filtration Rate, MRI – Magnetic Resonance Imaging, HFpEF – Heart Failure with Preserved Ejection Fraction, HFrEF – Heart Failure with Reduced Ejection Fraction, SV – Supraventricular, V – Ventricular LVEF – Left Ventricular Ejection Fraction, TAPSE – Tricuspid Annular Plane Systolic Excursion, PAPsys – Systolic Pulmonary Arterial Pressure, RV – Right Ventricle, LV – Left Ventricle, RA – Right Atrium, LA – Left Atrium, PPI – Proton Pump Inhibitor, NSAID – Nonsteroidal anti-inflammatory drug, RAAS – Renin-Angiotensin-Aldosterone System MMF – Mycophenolate mofetil, MTX – Methotrexate, CYC – Cyclophosphamide. Early SSc was defined as a disease duration of less than three years from the first non-Raynaud disease manifestation. SSc patients were classified as inflammatory if one or more of the following parameters were present at the time of biopsy: CRP > 0.6 mg/dl, thrombocyte count over 330,000/µl, arthritis, or myositis.

**Supplementary Table 2. Demographic and clinical data of the AMI, ICM, HCM and PVMF cohorts.**

| Cohort | Patient ID | Sex | Age (years) | Cardiotropic virus PCR | Source | Histopathology |
|---|---|---|---|---|---|---|
| PVMF | 1 | M | 43 | HHV6 | NA | Moderate interstitial fibrosis |
| PVMF | 2 | F | 38 | Parvovirus B19 | LV | Mild diffuse and focal interstitial fibrosis with focal perivascular accentuation |
| PVMF | 3 | F | 52 | HHV6 | LV | Moderate diffuse interstitial fibrosis with additional focal perivascular accentuation |
| PVMF | 4 | F | 22 | negative | LV | Mild diffuse interstitial fibrosis |
| PVMF | 5 | M | 59 | negative | LV | Moderate diffuse and focal interstitial fibrosis |
| PVMF | 6 | F | 53 | Parvovirus B19 | LV | Mild diffuse and focal interstitial fibrosis with focal perivascular accentuation |
| PVMF | 7 | M | 18 | negative | LV | Mild diffuse and focal interstitial fibrosis |

| Cohort | Patient ID | Sex | Age (years) | Cardiotropic virus PCR | Source | Histopathology |
|---|---|---|---|---|---|---|
| PVMF | 8 | M | 62 | negative | NA | Mild diffuse interstitial fibrosis with focal perivascular accentuation |
| ICM | 1 | F | 58 | NA | LV apex | Severe subendocardial and perivascular fibrosis |
| ICM | 2 | M | 66 | NA | LV apex | Severe subendocardial and interstitial fibrosis |
| ICM | 3 | M | 41 | NA | LV apex | Severe interstitial fibrosis |
| ICM | 4 | M | 76 | NA | LV apex | Moderate interstitial and subendocardial fibrosis |
| ICM | 5 | M | 58 | NA | LV apex | Severe interstitial fibrosis |
| ICM | 6 | M | 43 | NA | LV | Severe subendocardial and interstitial fibrosis |
| ICM | 7 | F | 44 | NA | LV | Severe interstitial fibrosis. Presence of perivascular fibrosis |
| ICM | 8 | M | 62 | NA | LV | Mild interstitial fibrosis |
| ICM | 9 | M | 43 | NA | LV | Severe subendocardial and interstitial fibrosis |
| HCM | 1 | M | 35 | NA | LV apex | Severe hypertrophy, focal interstitial fibrosis, mild to moderate subendocardial fibrosis |
| HCM | 2 | M | 57 | NA | LV apex | Severe hypertrophy, mild interstitial and subendocardial fibrosis |
| HCM | 3 | M | 32 | NA | LV apex | Mild hypertrophy, moderate interstitial fibrosis and mild subendocardial fibrosis |
| HCM | 4 | M | 4 | NA | LV | Severe hypertrophy and severe interstitial fibrosis |
| HCM | 5 | M | 21 | NA | LV | Severe hypertrophy, mild subendocardial and interstitial fibrosis |
| AMI | 1 | M | 51 | NA | LV apex | Extensive subacute, focally organizing myocardial infarction. Perivascular and subendocardial fibrosis. |
| AMI | 2 | M | 57 | NA | LV apex | Interstitial fibrosis and multiple acute myocardial infarcts showing neutrophilic demarcation. |

| Cohort | Patient ID | Sex | Age (years) | Cardiotropic virus PCR | Source | Histopathology |
|---|---|---|---|---|---|---|
| AMI | 3 | M | 61 | NA | LV apex | Coagulative necrosis, recent hemorrhage, and early leukocytic demarcation, consistent with a relatively recent acute myocardial infarction. |

F – female; M – male; NA – not available; PCR – polymerase chain reaction; HHV6 – human herpesvirus 6; LV – left ventricle.

**Supplementary Table 3. CosMx gene panel.**

| Panel entry | Panel entry | Panel entry | Panel entry | Panel entry | Panel entry | Panel entry |
|---|---|---|---|---|---|---|
| AATK | ABL1 | ABL2 | ACACB | ACE | ACKR1 | ACKR3 |
| ACKR4 | ACTA2 | ACTG2 | ACVR1 | ACVR1B | ACVR2A | ACVRL1 |
| ADGRA2 | ADGRA3 | ADGRE2 | ADGRE5 | ADGRF1 | ADGRF3 | ADGRF5 |
| ADGRG1 | ADGRG3 | ADGRG5 | ADGRG6 | ADGRL1 | ADGRL2 | ADGRL4 |
| ADGRV1 | ADIPOQ | ADIRF | ADM2 | AGR2 | AHI1 | AHR |
| AIF1 | AKT1 | ALCAM | ANGPT1 | ANGPT2 | ANGPTL1 | ANKRD1 |
| ANXA1 | ANXA2 | ANXA4 | **APCDD1** | APOA1 | APOC1 | APOD |
| APOE | APP | AQP3 | AR | AREG | ARF1 | ARG1 |
| ARHGDIB | ARID5B | ATF3 | ATG10 | ATG12 | ATG5 | ATM |
| ATP5F1E | ATR | AXL | AZGP1 | AZU1 | B2M | B3GNT7 |
| BAG3 | BAX | BBLN | BCL2 | BCL2L1 | BECN1 | BEST1 |
| BGN | BID | BIRC5 | BMP1 | BMP2 | BMP3 | BMP4 |
| BMP5 | BMP7 | BMPR1A | BMPR2 | BRCA1 | BST1 | BST2 |
| BTF3 | BTG1 | BTK | C11orf96 | C1QA | C1QB | C1QC |
| **C1QTNF3** | C5AR2 | CACNA1C | CALB1 | CALD1 | CALM1 | CALM2 |

| Panel entry | Panel entry | Panel entry | Panel entry | Panel entry | Panel entry | Panel entry |
|---|---|---|---|---|---|---|
| CALM3 | CAMP | CARMN | CASP3 | CASP8 | CASR | CAV1 |
| CCL11 | CCL13 | CCL15 | CCL18 | CCL19 | CCL2 | CCL20 |
| CCL21 | CCL26 | CCL28 | CCL3/L1/L3 | CCL4/L1/L2 | CCL5 | CCL8 |
| **CCN5** | CCND1 | CCR1 | CCR10 | CCR2 | CCR5 | CCR7 |
| CCRL2 | CD14 | CD163 | CD164 | CD19 | CD2 | CD209 |
| CD24 | CD27 | CD274 | CD276 | CD28 | CD300A | CD33 |
| CD34 | CD36 | CD37 | CD38 | CD3D | CD3E | CD3G |
| CD4 | CD40 | CD40LG | CD44 | CD47 | CD48 | CD52 |
| CD53 | CD55 | CD58 | CD59 | CD5L | CD63 | CD68 |
| CD69 | CD70 | CD74 | CD79A | CD80 | CD81 | CD83 |
| CD84 | CD86 | CD8A | CD8B | CD9 | CDH1 | CDH11 |
| CDH19 | CDH5 | CDKN1A | CDKN3 | CEACAM1 | CEACAM6 | CELSR1 |
| CELSR2 | CENPF | CFD | CFLAR | CHEK1 | CHEK2 | CHI3L1 |
| CIDEA | CIITA | CLCF1 | CLDN4 | CLEC10A | CLEC12A | CLEC14A |
| CLEC1A | CLEC2B | CLEC2D | CLEC4A | CLEC4D | CLEC4E | CLEC5A |
| CLEC7A | CLOCK | CLU | CMKLR1 | CNTFR | COL11A1 | COL12A1 |
| COL14A1 | COL15A1 | COL16A1 | COL17A1 | COL18A1 | COL1A1 | COL1A2 |
| COL21A1 | COL27A1 | COL3A1 | COL4A1 | COL4A2 | COL4A5 | COL5A1 |
| COL5A2 | COL5A3 | COL6A1 | COL6A2 | COL6A3 | COL8A1 | COL9A2 |
| COL9A3 | COTL1 | CPA3 | CPB1 | CRIP1 | CRP | CRYAB |
| CSF1 | CSF1R | CSF2 | CSF2RA | CSF2RB | CSF3 | CSF3R |
| CSHL1 | CSK | CST7 | **CTHRC1** | CTLA4 | CTNNB1 | CTSG |
| CTSW | CUZD1 | CX3CL1 | CX3CR1 | CXCL1/2/3 | CXCL10 | CXCL12 |
| CXCL14 | CXCL16 | CXCL17 | CXCL5 | CXCL8 | CXCL9 | CXCR1 |
| CXCR2 | CXCR3 | CXCR4 | CXCR5 | CXCR6 | CYP1B1 | CYP2U1 |

| **Panel entry** | **Panel entry** | **Panel entry** | **Panel entry** | **Panel entry** | **Panel entry** | **Panel entry** |
|---|---|---|---|---|---|---|
| CYSTM1 | CYTOR | DCN | DDC | DDIT3 | DDR1 | DDR2 |
| DDX58 | DHRS2 | DLL1 | DLL4 | DMBT1 | DNMT1 | DNMT3A |
| DPP4 | DST | DUSP1 | DUSP2 | DUSP4 | DUSP5 | DUSP6 |
| EFNA1 | EFNA4 | EFNA5 | EFNB1 | EFNB2 | EGF | EGFR |
| EIF5A/L1 | ELANE | EMP3 | ENG | ENO1 | ENTPD1 | EOMES |
| EPCAM | EPHA2 | EPHA3 | EPHA4 | EPHA7 | EPHB2 | EPHB3 |
| EPHB4 | EPHB6 | EPOR | ERBB2 | ERBB3 | ESAM | ESR1 |
| ETS1 | ETV4 | ETV5 | EZH2 | EZR | FABP4 | FABP5 |
| FAM30A | FAS | FASLG | FASN | FAU | **FBLN1** | FCER1G |
| FCGBP | FCGR3A/B | FCRLA | FES | FFAR2 | FFAR3 | FFAR4 |
| FGF1 | FGF12 | FGF13 | FGF18 | FGF2 | FGF7 | FGF9 |
| FGFR1 | FGFR2 | FGFR3 | FGG | FGR | FHIT | FKBP11 |
| FKBP5 | FLT1 | FLT3LG | FN1 | FOS | FOXF1 | FOXP3 |
| FPR1 | FYB1 | FYN | FZD1 | FZD3 | FZD4 | FZD5 |
| FZD6 | FZD7 | FZD8 | G6PD | GADD45B | GAS6 | GATA3 |
| GC | GCG | GDF15 | **GLI2** | GLUD1 | GLUL | GNLY |
| GPBAR1 | GPER1 | GPNMB | GPR183 | GPX1 | GPX3 | GSN |
| GSTP1 | GZMA | GZMB | GZMH | GZMK | H2AZ1 | H4C3 |
| HAVCR2 | HBA1/2 | HBB | HCAR2/3 | HCK | HCST | HDAC1 |
| HDAC11 | HDAC3 | HDAC4 | HDAC5 | HEY1 | HGF | HIF1A |
| HILPDA | HLA-DPA1 | HLA-DPB1 | HLA-DQA1 | HLA-DQB1/2 | HLA-DRA | HLA-DRB |
| HMGB2 | HMGN2 | HPGDS | HSD17B2 | HSP90AA1 | HSP90AB1 | HSP90B1 |
| HSPA1A/B | HSPB1 | HTT | IAPP | ICA1 | ICAM1 | ICAM2 |
| ICAM3 | ICOS | ICOSLG | IDO1 | IER3 | IFI27 | IFI44L |
| IFIH1 | IFIT1 | IFIT3 | IFITM1 | IFITM3 | IFNA1/13 | IFNAR1 |

| Panel entry | Panel entry | Panel entry | Panel entry | Panel entry | Panel entry | Panel entry |
|---|---|---|---|---|---|---|
| IFNAR2 | IFNG | IFNGR1 | IFNGR2 | IFNL2/3 | IGF1 | IGF1R |
| IGF2 | IGF2R | IGFBP3 | IGFBP5 | IGFBP6 | IGFBP7 | IGHA1 |
| IGHD | IGHG1 | IGHG2 | IGHM | IGKC | IKZF3 | IL10 |
| IL10RA | IL10RB | IL11 | IL11RA | IL12A | IL12B | IL12RB1 |
| IL12RB2 | IL13RA1 | IL15 | IL15RA | IL16 | IL17A | IL17B |
| IL17D | IL17RA | IL17RB | IL17RE | IL18 | IL18R1 | IL1A |
| IL1B | IL1R1 | IL1R2 | IL1RAP | IL1RL1 | IL1RN | IL2 |
| IL20 | IL20RA | IL22RA1 | IL23A | IL24 | IL27RA | IL2RA |
| IL2RB | IL2RG | IL32 | IL33 | IL34 | IL36G | IL3RA |
| IL4R | IL6 | IL6R | IL6ST | IL7 | IL7R | INHA |
| INHBA | INHBB | INS | INSR | IRF3 | IRF4 | ITGA1 |
| ITGA2 | ITGA3 | ITGA5 | ITGA6 | ITGA8 | ITGA9 | ITGAE |
| ITGAL | ITGAM | ITGAV | ITGAX | ITGB1 | ITGB2 | ITGB4 |
| ITGB5 | ITGB6 | ITGB8 | ITK | ITM2A | JAG1 | JAK1 |
| JAK2 | JCHAIN | JUN | JUNB | KDR | KIT | KITLG |
| KLF2 | KLK3 | KLRB1 | KLRF1 | KLRK1 | KRAS | KRT1 |
| KRT10 | KRT13 | KRT14 | KRT15 | KRT16 | KRT17 | KRT18 |
| KRT19 | KRT20 | KRT23 | KRT4 | KRT5 | KRT6A/B/C | KRT7 |
| KRT8 | KRT80 | KRT86 | LAG3 | LAIR1 | LAMP2 | LAMP3 |
| LCN2 | LDB2 | LDHA | LDLR | LEFTY1 | LEP | LGALS1 |
| LGALS3 | LGALS3BP | LGALS9 | LGR5 | LIF | LIFR | LINC01781 |
| LINC01857 | LINC02446 | LMNA | LPAR5 | **LRRC15** | LTB | LTBR |
| LTF | LUM | LY6D | LY75 | LYN | LYZ | MAF |
| MALAT1 | MAML2 | MAP1LC3B/2 | MAPK13 | MAPK14 | MARCO | MB |
| MECOM | MEG3 | MERTK | MET | MFAP5 | MGP | MHC I |

| Panel entry | Panel entry | Panel entry | Panel entry | Panel entry | Panel entry | Panel entry |
|---|---|---|---|---|---|---|
| MIF | MIR4435-2HG | MKI67 | MMP1 | MMP12 | MMP14 | MMP19 |
| MMP2 | MMP7 | MMP9 | MPO | MRC1 | MRC2 | MS4A1 |
| MS4A4A | MS4A6A | MSMB | MSR1 | MST1R | **MT-CO3** | MT1X |
| MT2A | MTOR | MX1 | MXRA8 | MYC | MYH11 | MYH6 |
| MYL12A | MYL4 | MYL7 | MYL9 | MZB1 | MZT2A/B | NACA |
| NANOG | **NBL1** | NCR1 | NDRG1 | NEAT1 | NELL2 | NFKB1 |
| NFKBIA | NGFR | NKG7 | NLRC4 | NLRC5 | NLRP1 | NLRP2 |
| NLRP3 | NOD2 | NOSIP | NOTCH1 | NOTCH2 | NOTCH3 | NPPC |
| NPR1 | NPR2 | NPR3 | NR1H2 | NR1H3 | NR3C1 | NRG1 |
| NRXN1 | NRXN3 | NTRK2 | NUSAP1 | OAS1 | OAS2 | OAS3 |
| OASL | OLFM4 | OLR1 | OSM | OSMR | P2RX5 | PARP1 |
| PCNA | PDCD1 | PDCD1LG2 | PDGFA | PDGFB | PDGFC | PDGFD |
| PDGFRA | PDGFRB | PDS5A | PECAM1 | PF4/V1 | PFN1 | PGF |
| PGR | PHLDA2 | **PI16** | **PIEZO1** | PIGR | PLAC8 | PLAC9 |
| PNOC | **POSTN** | POU5F1 | PPARA | PPARD | PPARG | PPIA |
| PRF1 | PRSS2 | PRTN3 | PSAP | PSCA | PSD3 | PTGDR2 |
| PTGDS | PTGES | PTGES2 | PTGES3 | PTGIS | PTGS1 | PTGS2 |
| PTK2 | PTK6 | PTPRC | PTPRCAP | PTTG1 | QRFPR | RAC1 |
| RAC2 | RACK1 | RAG1 | RAMP1 | RAMP2 | RAMP3 | RARA |
| RARB | RARG | RARRES1 | RARRES2 | RB1 | RBM47 | RBPJ |
| REG1A | RELA | RELT | RGCC | RGS1 | RGS2 | RGS5 |
| RNF43 | ROR1 | RORA | RPL21 | RPL22 | RPL32 | RPL34 |
| RPL37 | **RPS27** | RPS4Y1 | RSPO3 | RUNX3 | RXRA | RXRB |
| RYK | RYR2 | S100A10 | S100A2 | S100A4 | S100A6 | S100A8 |
| S100A9 | S100B | S100P | SAA1/2 | SAT1 | **SCARA5** | SCG5 |

| Panel entry | Panel entry | Panel entry | Panel entry | Panel entry | Panel entry | Panel entry |
|---|---|---|---|---|---|---|
| SCGB3A1 | SEC23A | SEC61G | SELENOP | SELL | SELPLG | SERPINA1 |
| SERPINA3 | SERPINB5 | SERPINH1 | SFN | **SFRP2** | SH3BGRL3 | SIGIRR |
| SLC2A1 | SLC40A1 | SLPI | SMAD2 | SMAD3 | SMAD4 | SMARCB1 |
| SMO | SNAI1 | SNAI2 | SOD1 | SOD2 | SORBS1 | SOSTDC1 |
| SOX2 | SOX4 | SOX9 | **SPARC** | SPARCL1 | SPINK1 | SPOCK2 |
| SPP1 | SPRY2 | SPRY4 | SQSTM1 | SRC | SREBF1 | SRGN |
| SST | ST6GAL1 | ST6GALNAC3 | STAT1 | STAT3 | STAT4 | STAT5A |
| STAT5B | STAT6 | **STC2** | STMN1 | SYK | TACSTD2 | TAGLN |
| TAP1 | TAP2 | TBX21 | TCAP | TCF7 | TCL1A | TEK |
| TFEB | TGFB1 | TGFB2 | TGFB3 | TGFBR1 | TGFBR2 | THBS1 |
| THBS2 | THSD4 | TIE1 | TIGIT | TIMP1 | TLR1 | TLR2 |
| TLR3 | TLR4 | TLR5 | TLR7 | TLR8 | TM4SF1 | TNF |
| TNFAIP6 | TNFRSF10A | TNFRSF10B | TNFRSF10D | TNFRSF11A | TNFRSF11B | TNFRSF12A |
| TNFRSF13B | TNFRSF14 | TNFRSF17 | TNFRSF18 | TNFRSF19 | TNFRSF1A | TNFRSF1B |
| TNFRSF21 | TNFRSF4 | TNFRSF9 | TNFSF10 | TNFSF12 | TNFSF13B | TNFSF14 |
| TNFSF15 | TNFSF4 | TNFSF8 | TNFSF9 | **TNMD** | TNNC1 | TNNT2 |
| TNXA/B | TOP2A | TOX | TP53 | TPM1 | TPM2 | **TPRA1** |
| TPSAB1/B2 | TPT1 | TSC22D1 | TSHZ2 | TTN | TTR | TUBB |
| TUBB4B | TWIST1 | TWIST2 | TXK | TYK2 | TYMS | TYROBP |
| UBA52 | UBE2C | UPK3A | VCAM1 | VCAN | VEGFA | VEGFB |
| VEGFC | VEGFD | VHL | VIM | VPREB3 | VSIR | VTN |
| VWA1 | VWF | WIF1 | WNT10B | WNT11 | WNT3 | WNT5A |
| WNT5B | WNT7A | WNT7B | WNT9A | XBP1 | XCL1/2 | XKR4 |
| YBX3 | YES1 | ZFP36 | | | | |

| Panel entry | Panel entry | Panel entry | Panel entry | Panel entry | Panel entry | Panel entry |
|---|---|---|---|---|---|---|
| **Negative controls (n = 10)** | | | | | | |
| Negative01 | Negative02 | Negative03 | Negative04 | Negative05 | Negative06 | Negative07 |
| Negative08 | Negative09 | Negative10 | | | | |

The panel comprised 950 core gene targets, 19 custom gene targets, and 10 negative controls (979 entries in total). **Custom additions are shown in bold**; core targets are in regular type.

#### Supplementary Table 4. CODEX panel

| Target | Clone | Manufacturer | Catalog number | Fluorophore | Dilution (1:X) |
|---|---|---|---|---|---|
| POSTN | Polyclonal | Abcam | ab14041 | ATTO550 | 150 |
| TNNT2 | 1C11 | Abcam | ab8295 | ATTO647 | 50 |
| GSN | Polyclonal | Proteintech | 11644-2-AP | ATTO550 | 150 |
| JUNB | Polyclonal | Proteintech | 10486-1-AP | ATTO647 | 150 |
| NOTCH3 | Polyclonal | Proteintech | 55114-1-AP | ATTO550 | 150 |
| TIMP1 | Polyclonal | Proteintech | 16644-1-AP | ATTO647 | 150 |
| C1QC | Polyclonal | Proteintech | 16889-1-AP | ATTO550 | 150 |
| APOE | Polyclonal | Proteintech | 18254-1-AP | ATTO647 | 300 |

| | | | | | |
|---|---|---|---|---|---|
| SPP1 | Polyclonal | Proteintech | 22952-1-AP | ATTO550 | 50 |
| DCN | 115402 | R&D | MAB143 | ATTO647 | 150 |
| CD68 | KP-1 | Biolegend | 916104 | ATTO550 | 50 |
| CD45 | 2B11 + PD7/26 | Novus | NBP2-34528 | ATTO647 | 300 |
| αSMA | 1A4 | Biolegend | 614852 | ATTO550 | 150 |
| COL1 | EPR7785 | Abcam | Ab138492 | ATTO647 | 150 |
| CD31 | C31.3+C31.7+C31.10 | Novus Bio | NBP2-47785 | ATTO550 | 300 |
| FN1 | FN3 | eBioscience | 14-9869-82 | ATTO647 | 150 |
| vWF | 2F2-A9 (RUO) | BD | 555849 | ATTO550 | 100 |

#### Supplementary Figures and Figure Legends

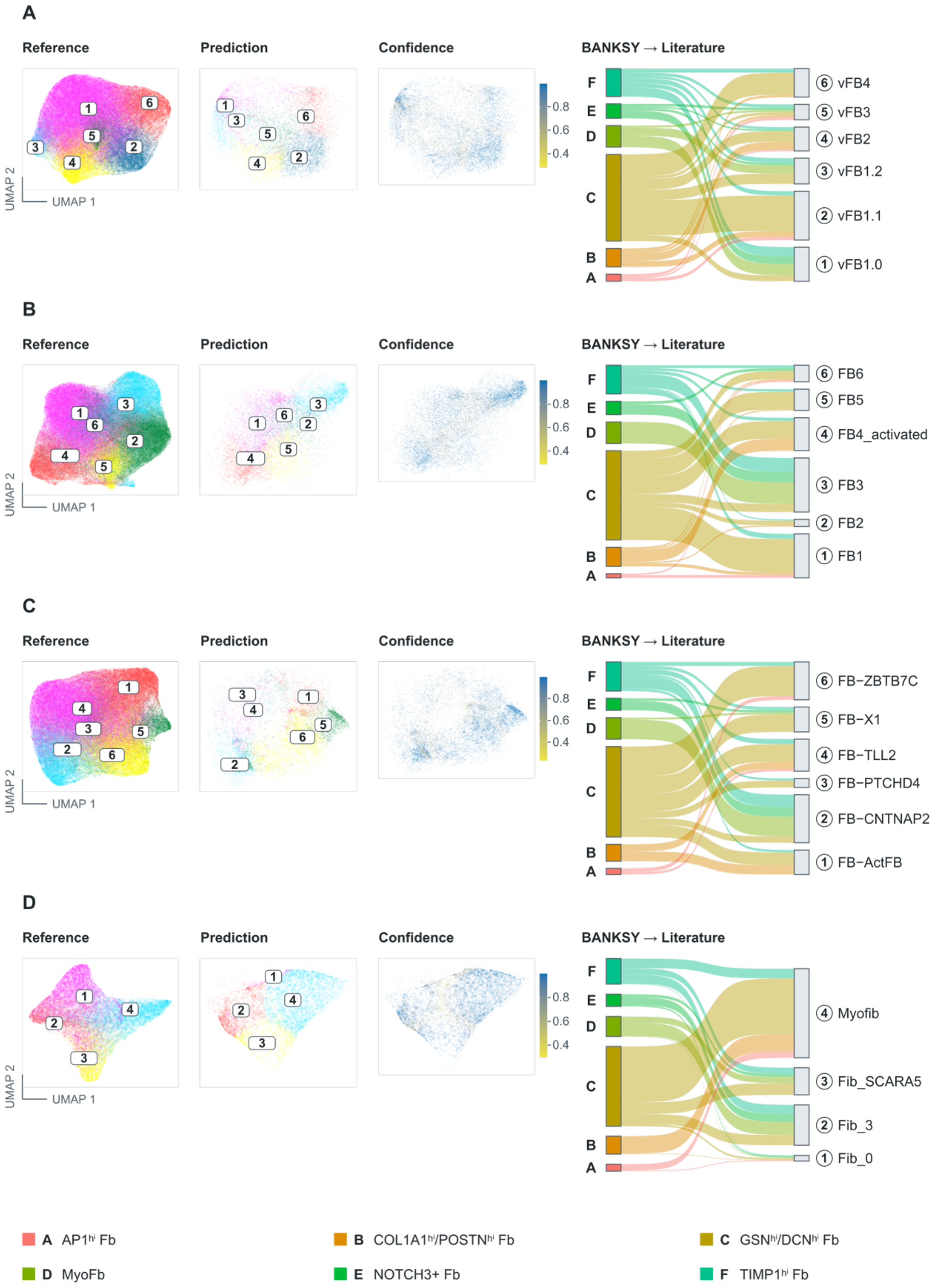
A
Reference
Prediction
Confidence
BANKSY → Literature
UMAP 1
UMAP 2
0.8
0.6
0.4
F
E
D
C
B
A
6 vFB4
5 vFB3
4 vFB2
3 vFB1.2
2 vFB1.1
1 vFB1.0
B
Reference
Prediction
Confidence
BANKSY → Literature
6 FB6
5 FB5
4 FB4_activated
3 FB3
2 FB2
1 FB1
C
Reference
Prediction
Confidence
BANKSY → Literature
6 FB−ZBTB7C
5 FB−X1
4 FB−TLL2
3 FB−PTCHD4
2 FB−CNTNAP2
1 FB−ActFB
D
Reference
Prediction
Confidence
BANKSY → Literature
4 Myofib
3 Fib_SCARA5
2 Fib_3
1 Fib_0
A AP1$^{hi}$ Fb
B COL1A1$^{hi}$/POSTN$^{hi}$ Fb
C GSN$^{hi}$/DCN$^{hi}$ Fb
D MyoFb
E NOTCH3+ Fb
F TIMP1$^{hi}$ Fb

**Supplementary Figure 1: Cross-dataset integration of fibroblast states with published reference atlases using scANVI**. Joint latent space of the query fibroblast dataset and reference atlas, shown as UMAP projections colored by reference annotation, scANVI-predicted fibroblast subtype, and prediction probability. Prediction probability reflects label-transfer confidence. Sankey plots showing correspondence between BANKSY-defined fibroblast clusters in the present dataset and fibroblast states in published reference atlases [1,3,29,30]. Flow width indicates the proportion of cells assigned to each reference subtype. UMAP, uniform manifold approximation and projection. Reference datasets were: **(A)** Reichart et al., European Genome-phenome Archive study EGAS00001006374, dataset EGAD00001009292; **(B)** Litviňuková et al., Human Cell Atlas/European Nucleotide Archive accession ERP123138; **(C)** Chaffin et al., Broad Institute Single Cell Portal project SCP1303; and **(D)** Kuppe et al., Human Cell Atlas project e9f36305-d857-44a3-93f0-df4e6007dc97, with processed data additionally available through CELLxGENE collection 8191c283-0816-424b-9b61-c3e1d6258a77 and Zenodo record 6578047.

**A.** **B.**

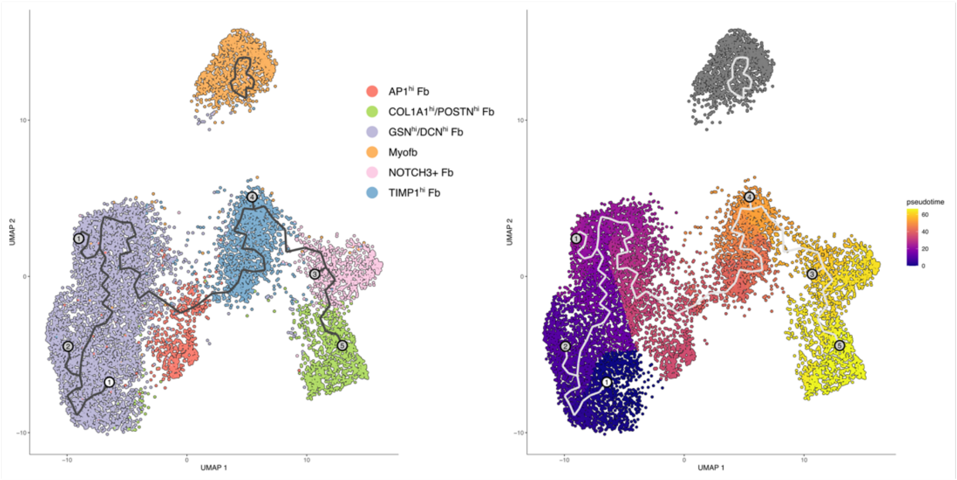

AP1hi Fb
COL1A1hi/POSTNhi Fb
GSNhi/DCNhi Fb
Myofb
NOTCH3+ Fb
TIMP1hi Fb
pseudotime
UMAP 1
UMAP 2


**C.**

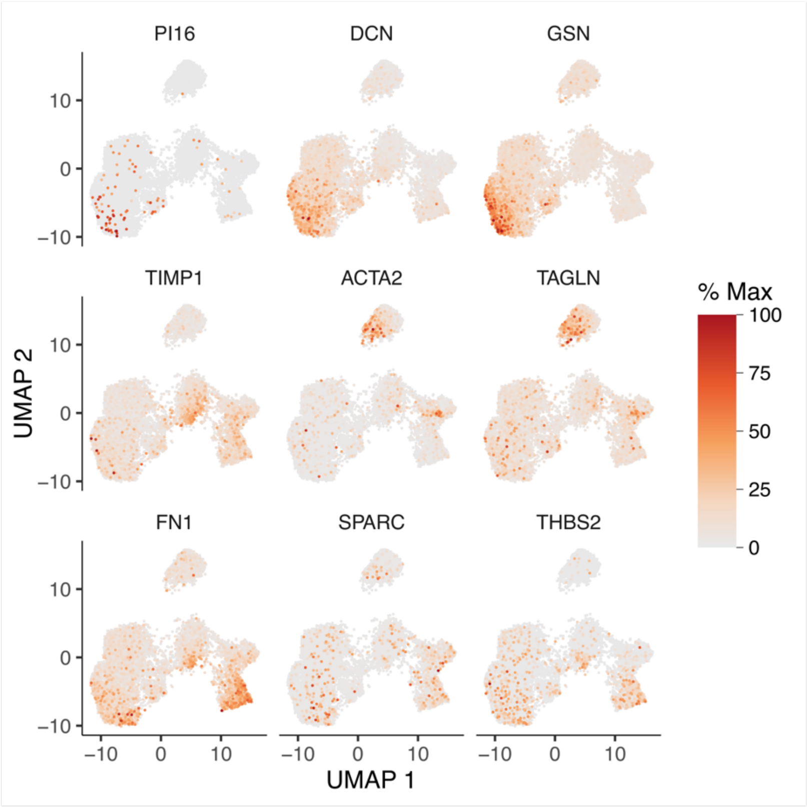

PI16
DCN
GSN
TIMP1
ACTA2
TAGLN
FN1
SPARC
THBS2
% Max
100
75
50
25
0
UMAP 2
UMAP 1

**Supplementary Figure 2**: **Pseudotemporal trajectory of fibroblast activation inferred by Monocle. (A)** Monocle-based trajectory analysis of fibroblast subpopulations, rooted in the $GSN^{hi}/DCN^{hi}$ Fb state and projected onto the UMAP embedding. The numbered circles denote principal graph nodes along the inferred trajectory. **(B)** Pseudotime coloring of the same UMAP reveals a continuum from early, more quiescent fibroblast states to later, more activated and differentiated states. **(C)** Feature plots showing the distribution of selected genes along the trajectory. Gene expression is shown scaled to each gene's maximum across the dataset. UMAP, uniform manifold approximation and projection.

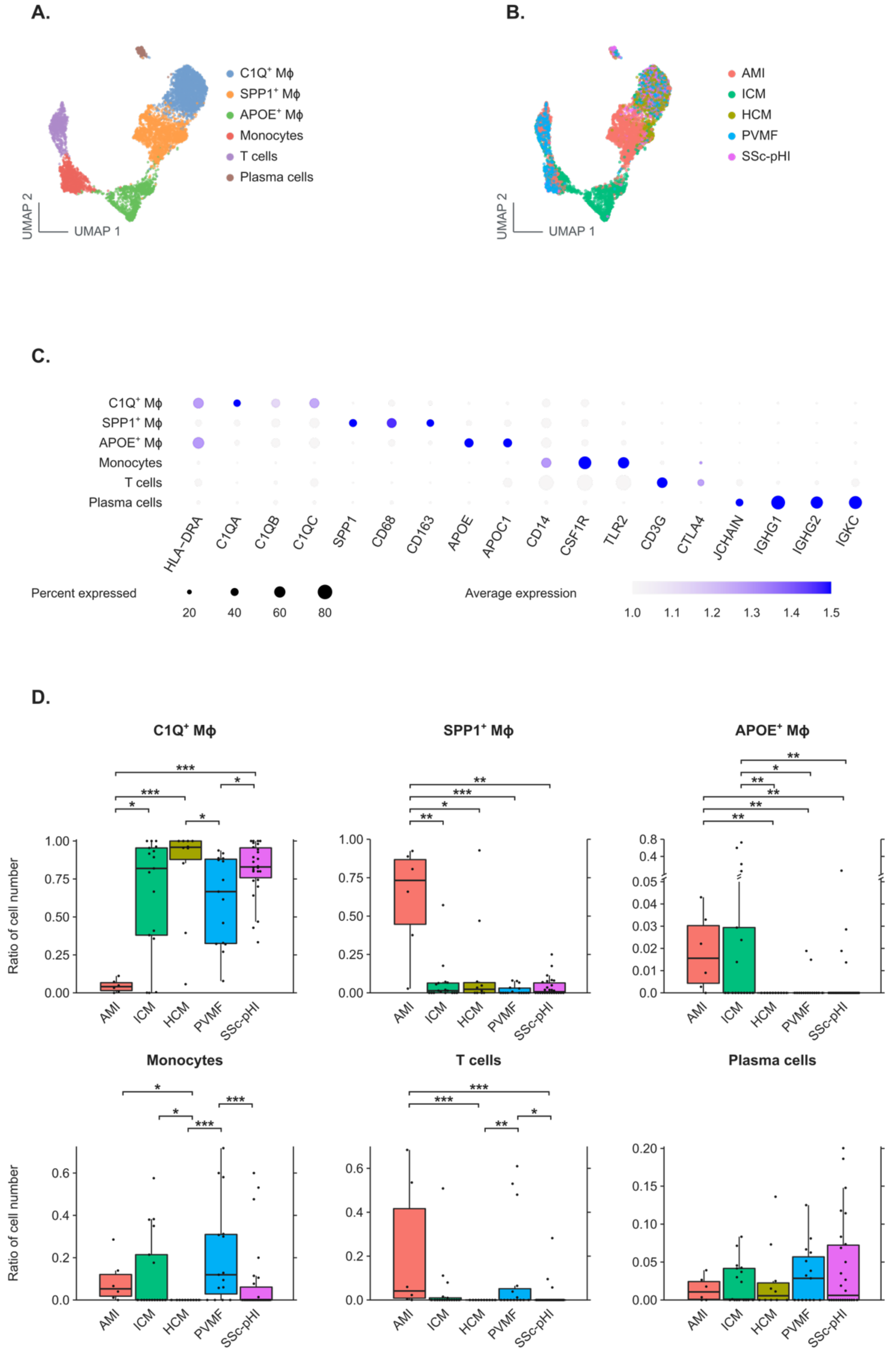
A.
C1Q⁺ Mϕ
SPP1⁺ Mϕ
APOE⁺ Mϕ
Monocytes
T cells
Plasma cells
UMAP 2
UMAP 1
B.
AMI
ICM
HCM
PVMF
SSc-pHI
UMAP 2
UMAP 1
C.
C1Q⁺ Mϕ
SPP1⁺ Mϕ
APOE⁺ Mϕ
Monocytes
T cells
Plasma cells
HLA-DRA
C1QA
C1QB
C1QC
SPP1
CD68
CD163
APOE
APOC1
CD14
CSF1R
TLR2
CD3G
CTLA4
JCHAIN
IGHG1
IGHG2
IGKC
Percent expressed
20
40
60
80
Average expression
1.0
1.1
1.2
1.3
1.4
1.5
D.
C1Q⁺ Mϕ
SPP1⁺ Mϕ
APOE⁺ Mϕ
Ratio of cell number
Monocytes
T cells
Plasma cells
Ratio of cell number
AMI
ICM
HCM
PVMF
SSc-pHI

**Supplementary Figure 3. Immune cell subcluster composition and distribution across disease cohorts. (A)** UMAP visualization of immune cells colored by annotated cell subclusters. **(B)** UMAP representation of the same immune-cell populations colored according to disease cohort (AMI, ICM, HCM, PVMF, and SSc-pHI), illustrating the distribution of cells from each disease condition across immune subclusters. **(C)** Dot plot showing expression of selected canonical marker genes. Dot size represents the percentage of cells expressing each gene, and color intensity represents the average expression level. **(D)** Box plots comparing the relative abundance of $C1QC^+$ MΦ, $SPP1^+$ MΦ, $APOE^+$ MΦ, monocytes, T cells, and plasma cells across disease conditions. Each dot represents an individual sample. Statistical significance (Kruskal-Wallis test) between groups is indicated as *$p < 0.05$, **$p < 0.01$, and ***$p < 0.001$. MΦ, macrophages; UMAP, uniform manifold approximation and projection.

**A.**

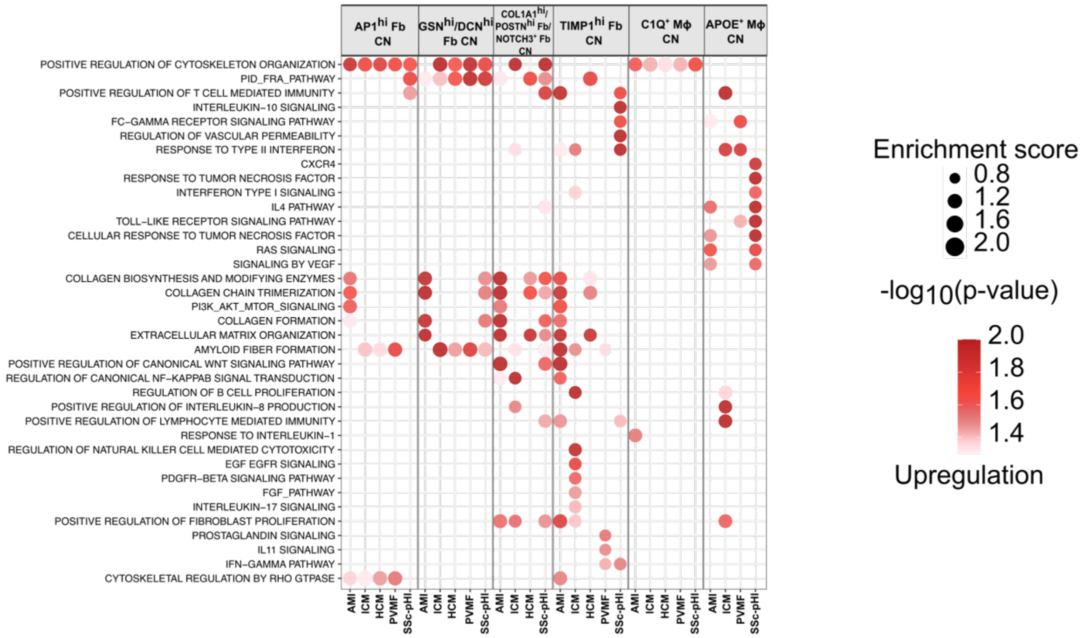

AP1hi Fb CN
GSNhi/DCNhi Fb CN
COL1A1hi/POSTNhi Fb/NOTCH3+ Fb CN
TIMP1hi Fb CN
C1Q+ Mϕ CN
APOE+ Mϕ CN
POSITIVE REGULATION OF CYTOSKELETON ORGANIZATION
PID_FRA_PATHWAY
POSITIVE REGULATION OF T CELL MEDIATED IMMUNITY
INTERLEUKIN–10 SIGNALING
FC–GAMMA RECEPTOR SIGNALING PATHWAY
REGULATION OF VASCULAR PERMEABILITY
RESPONSE TO TYPE II INTERFERON
CXCR4
RESPONSE TO TUMOR NECROSIS FACTOR
INTERFERON TYPE I SIGNALING
IL4 PATHWAY
TOLL–LIKE RECEPTOR SIGNALING PATHWAY
CELLULAR RESPONSE TO TUMOR NECROSIS FACTOR
RAS SIGNALING
SIGNALING BY VEGF
COLLAGEN BIOSYNTHESIS AND MODIFYING ENZYMES
COLLAGEN CHAIN TRIMERIZATION
PI3K_AKT_MTOR_SIGNALING
COLLAGEN FORMATION
EXTRACELLULAR MATRIX ORGANIZATION
AMYLOID FIBER FORMATION
POSITIVE REGULATION OF CANONICAL WNT SIGNALING PATHWAY
REGULATION OF CANONICAL NF–KAPPAB SIGNAL TRANSDUCTION
REGULATION OF B CELL PROLIFERATION
POSITIVE REGULATION OF INTERLEUKIN–8 PRODUCTION
POSITIVE REGULATION OF LYMPHOCYTE MEDIATED IMMUNITY
RESPONSE TO INTERLEUKIN–1
REGULATION OF NATURAL KILLER CELL MEDIATED CYTOTOXICITY
EGF EGFR SIGNALING
PDGFR–BETA SIGNALING PATHWAY
FGF_PATHWAY
INTERLEUKIN–17 SIGNALING
POSITIVE REGULATION OF FIBROBLAST PROLIFERATION
PROSTAGLANDIN SIGNALING
IL11 SIGNALING
IFN–GAMMA PATHWAY
CYTOSKELETAL REGULATION BY RHO GTPASE
AMI
ICM
HCM
PVMF
SSc-pHI
Enrichment score
0.8
1.2
1.6
2.0
-log10(p-value)
2.0
1.8
1.6
1.4
Upregulation


**B.**

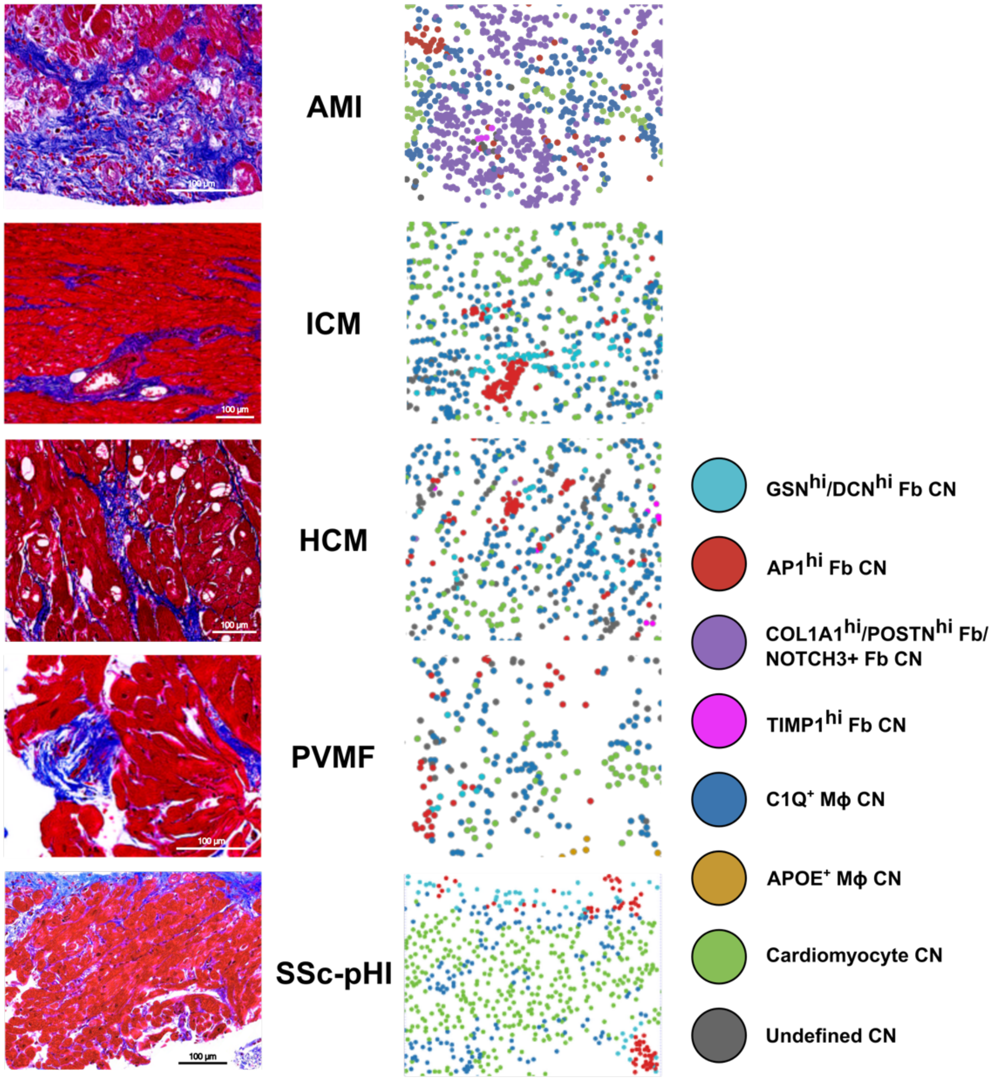

AMI
ICM
HCM
PVMF
SSc-pHI
100 µm
GSNhi/DCNhi Fb CN
AP1hi Fb CN
COL1A1hi/POSTNhi Fb/
NOTCH3+ Fb CN
TIMP1hi Fb CN
C1Q+ Mϕ CN
APOE+ Mϕ CN
Cardiomyocyte CN
Undefined CN

**Supplementary Figure 4: Pathway enrichment and spatial localization of cellular neighborhoods across disease conditions. (A)** Dot plot of selected pathway enrichment scores across fibroblast populations under different disease conditions. Enrichment analysis was performed using fgsea on pseudo-bulk expression matrices generated by averaging expression within each fibroblast subpopulation for each disease condition and implemented with the decoupleR R package. Pathways with $p < 0.05$ were retained. Statistical significance was assessed by comparing estimated marginal means from linear mixed-effects models including donor-specific random intercepts. Dot size indicates enrichment score, and color intensity represents normalized $-\log10(p)$ values for upregulated and downregulated pathways. **(B)** Representative images illustrating the spatial localization of cellular neighborhoods in myocardial biopsy samples. Left, Masson's trichrome-stained tissue sections, scale bar for reference. Right, corresponding spatial maps showing the distribution of annotated cellular neighborhoods within the same biopsy specimens. Colors denote individual cellular neighborhoods as indicated.

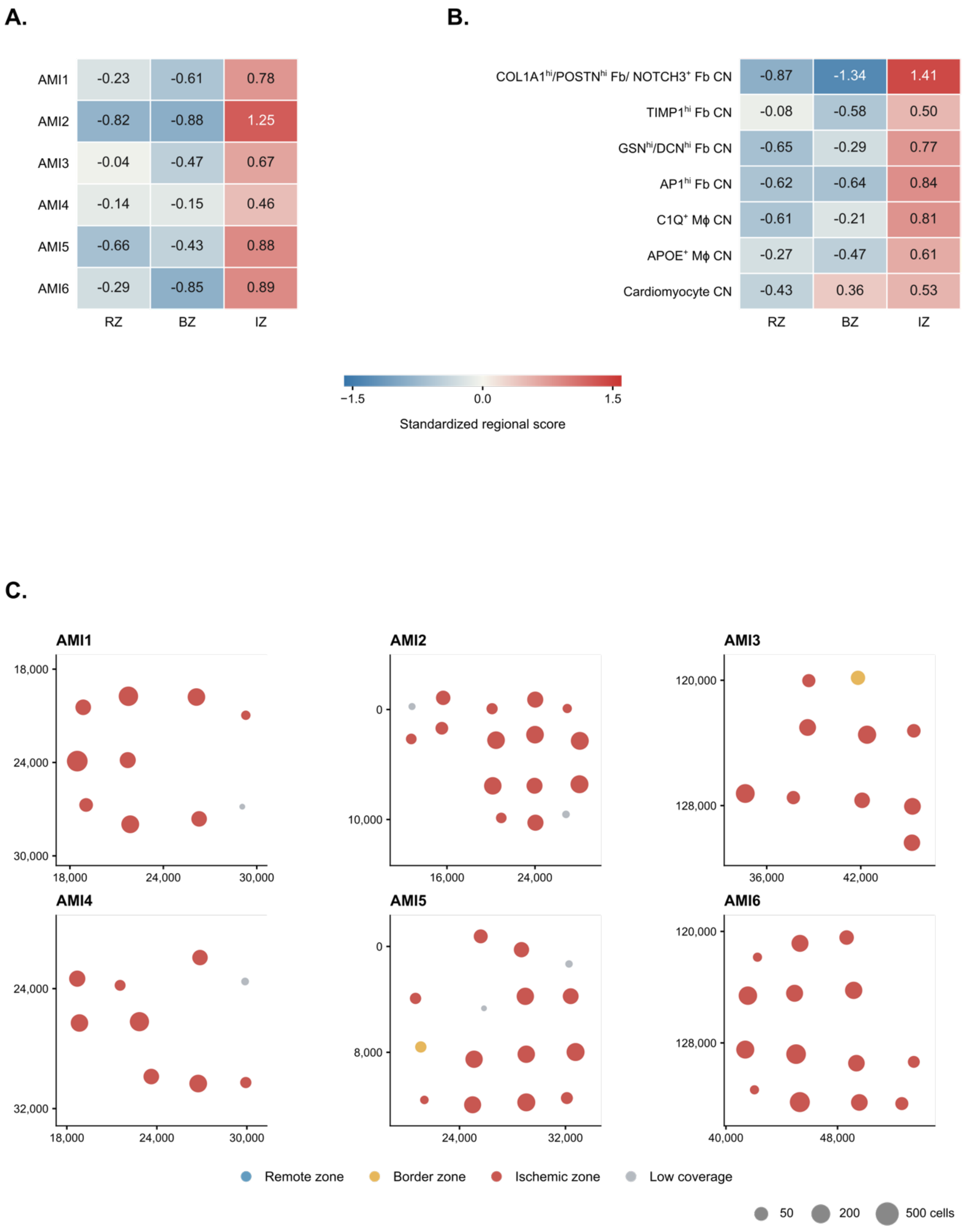

A.
AMI1 -0.23 -0.61 0.78
AMI2 -0.82 -0.88 1.25
AMI3 -0.04 -0.47 0.67
AMI4 -0.14 -0.15 0.46
AMI5 -0.66 -0.43 0.88
AMI6 -0.29 -0.85 0.89
RZ BZ IZ
B.
COL1A1hi/POSTNhi Fb/ NOTCH3+ Fb CN -0.87 -1.34 1.41
TIMP1hi Fb CN -0.08 -0.58 0.50
GSNhi/DCNhi Fb CN -0.65 -0.29 0.77
AP1hi Fb CN -0.62 -0.64 0.84
C1Q+ Mϕ CN -0.61 -0.21 0.81
APOE+ Mϕ CN -0.27 -0.47 0.61
Cardiomyocyte CN -0.43 0.36 0.53
RZ BZ IZ
−1.5 0.0 1.5
Standardized regional score
C.
AMI1
AMI2
AMI3
AMI4
AMI5
AMI6
Remote zone
Border zone
Ischemic zone
Low coverage
50
200
500 cells

**Supplementary Figure 5: Transcriptomic correspondence of acute myocardial infarction and cellular neighborhoods to infarction regions.** **(A)** Standardized regional scores for acute myocardial infarction samples relative to the remote-zone (RZ), border-zone (BZ), and ischemic-zone (IZ) signatures derived from Kuppe et al. [3]. **(B)** Mean regional scores for seven cellular neighborhoods (CNs), with equal weighting of eligible samples. **(C)** ROI correspondence maps for the six AMI samples (AMI1-AMI6). Dots mark the mean native coordinates of retained cells within each ROI. Colors indicate the highest-scoring reference region; grey denotes insufficient coverage. Dot area scales with the square root of cell number.

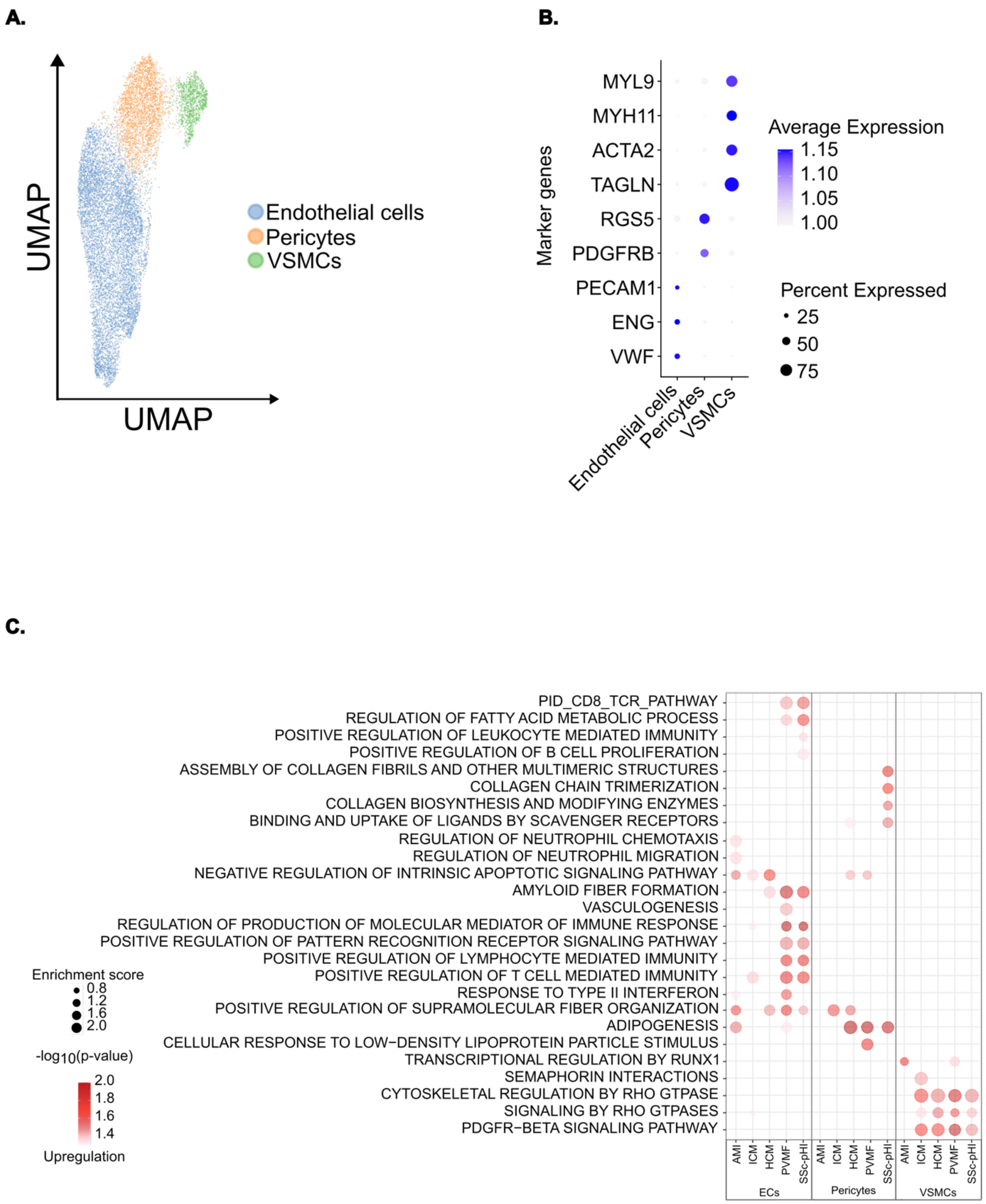
A.
UMAP
UMAP
Endothelial cells
Pericytes
VSMCs
B.
Marker genes
MYL9
MYH11
ACTA2
TAGLN
RGS5
PDGFRB
PECAM1
ENG
VWF
Endothelial cells
Pericytes
VSMCs
Average Expression
1.15
1.10
1.05
1.00
Percent Expressed
25
50
75
C.
PID_CD8_TCR_PATHWAY
REGULATION OF FATTY ACID METABOLIC PROCESS
POSITIVE REGULATION OF LEUKOCYTE MEDIATED IMMUNITY
POSITIVE REGULATION OF B CELL PROLIFERATION
ASSEMBLY OF COLLAGEN FIBRILS AND OTHER MULTIMERIC STRUCTURES
COLLAGEN CHAIN TRIMERIZATION
COLLAGEN BIOSYNTHESIS AND MODIFYING ENZYMES
BINDING AND UPTAKE OF LIGANDS BY SCAVENGER RECEPTORS
REGULATION OF NEUTROPHIL CHEMOTAXIS
REGULATION OF NEUTROPHIL MIGRATION
NEGATIVE REGULATION OF INTRINSIC APOPTOTIC SIGNALING PATHWAY
AMYLOID FIBER FORMATION
VASCULOGENESIS
REGULATION OF PRODUCTION OF MOLECULAR MEDIATOR OF IMMUNE RESPONSE
POSITIVE REGULATION OF PATTERN RECOGNITION RECEPTOR SIGNALING PATHWAY
POSITIVE REGULATION OF LYMPHOCYTE MEDIATED IMMUNITY
POSITIVE REGULATION OF T CELL MEDIATED IMMUNITY
RESPONSE TO TYPE II INTERFERON
POSITIVE REGULATION OF SUPRAMOLECULAR FIBER ORGANIZATION
ADIPOGENESIS
CELLULAR RESPONSE TO LOW−DENSITY LIPOPROTEIN PARTICLE STIMULUS
TRANSCRIPTIONAL REGULATION BY RUNX1
SEMAPHORIN INTERACTIONS
CYTOSKELETAL REGULATION BY RHO GTPASE
SIGNALING BY RHO GTPASES
PDGFR−BETA SIGNALING PATHWAY
AMI
ICM
HCM
PVMF
SSc-pHI
ECs
AMI
ICM
HCM
PVMF
SSc-pHI
Pericytes
AMI
ICM
HCM
PVMF
SSc-pHI
VSMCs
Enrichment score
0.8
1.2
1.6
2.0
-log10(p-value)
2.0
1.8
1.6
1.4
Upregulation

**Supplementary Figure 6**: **Vascular cell subsets and their functions across disease subtypes.** **(A)** UMAP embedding of vascular cells showing the major vascular cell populations identified by BANKSY clustering. **(B)** Dot plot of marker genes used for vascular cell annotation. Dot size indicates the proportion of cells expressing a given gene, and color intensity represents average expression level. **(C)** Dot plot of selected pathway enrichment scores across vascular populations under different disease conditions. Enrichment analysis was performed using fgsea on pseudo-bulk expression matrices generated by averaging expression within each vascular cell subpopulation for each disease condition and implemented through the decoupleR R package. Pathways with $p < 0.05$ were retained. Statistical significance was assessed by comparing estimated marginal means from linear mixed-effects models including donor-specific random intercepts. Dot size indicates enrichment score, and color intensity represents normalized $-\log10(p)$ values for upregulation. UMAP, uniform manifold approximation and projection.

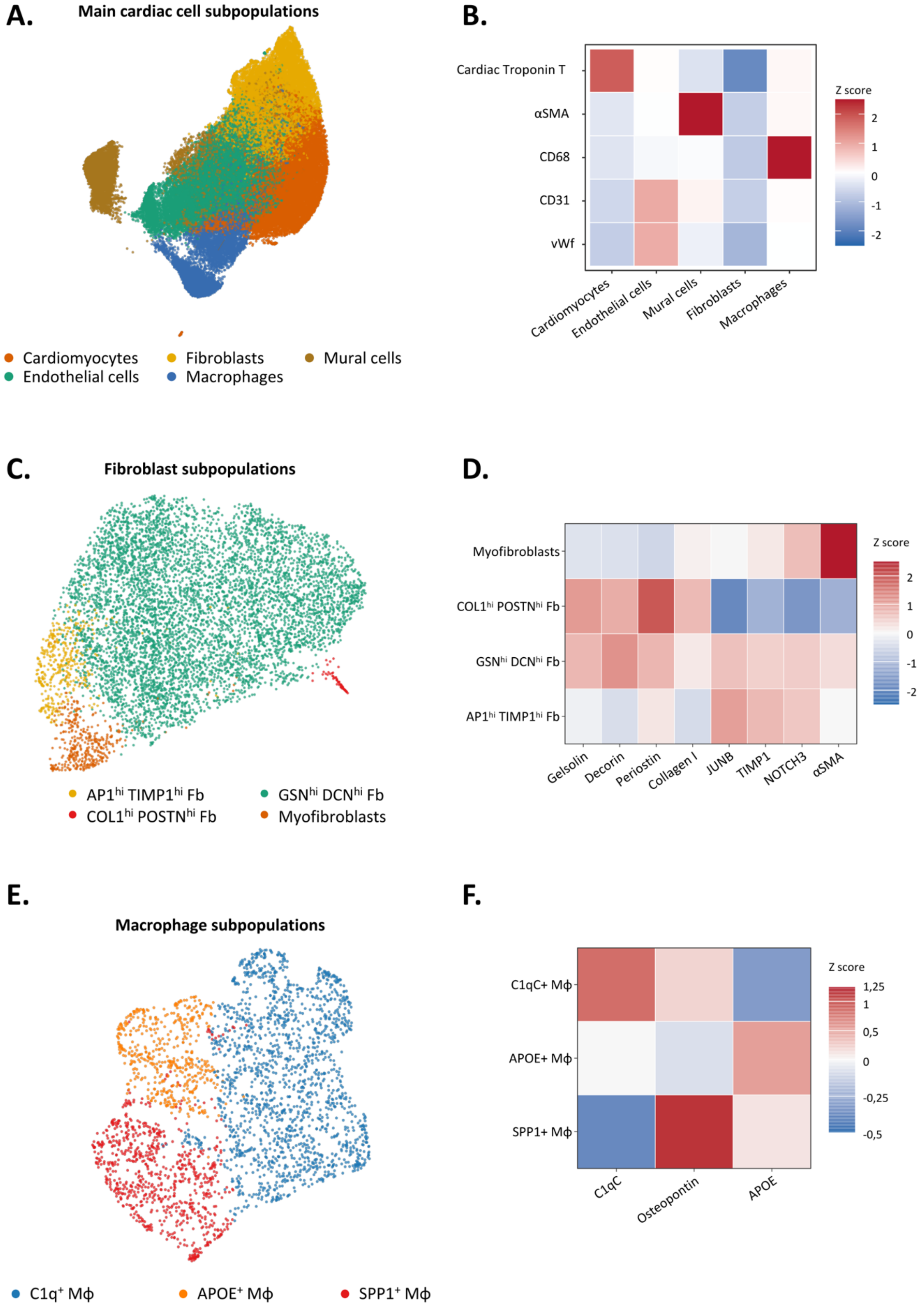
A.
Main cardiac cell subpopulations
Cardiomyocytes
Fibroblasts
Mural cells
Endothelial cells
Macrophages
B.
Cardiac Troponin T
αSMA
CD68
CD31
vWf
Z score
2
1
0
-1
-2
Cardiomyocytes
Endothelial cells
Mural cells
Fibroblasts
Macrophages
C.
Fibroblast subpopulations
AP1hi TIMP1hi Fb
GSNhi DCNhi Fb
COL1hi POSTNhi Fb
Myofibroblasts
D.
Myofibroblasts
COL1hi POSTNhi Fb
GSNhi DCNhi Fb
AP1hi TIMP1hi Fb
Z score
2
1
0
-1
-2
Gelsolin
Decorin
Periostin
Collagen I
JUNB
TIMP1
NOTCH3
αSMA
E.
Macrophage subpopulations
C1q+ Mϕ
APOE+ Mϕ
SPP1+ Mϕ
F.
C1qC+ Mϕ
APOE+ Mϕ
SPP1+ Mϕ
Z score
1,25
1
0,5
0
-0,25
-0,5
C1qC
Osteopontin
APOE

**Supplementary Figure 7: Marker-based annotation of main cardiac cell populations and fibroblast and macrophage subpopulations in CODEX.** **(A)** UMAP coloured according to the main cardiac cell populations. Each point represents one cell. **(B)** Heatmap showing the protein marker expression of the major cell populations. Rows indicate markers and columns indicate annotated cell subsets. Colours represent within-marker robust *z* scores, with red and blue indicating higher and lower relative abundance, respectively. **(C, E)** UMAPs coloured according to subpopulations of fibroblasts **(C)** or macrophages **(E)**, respectively. **(D, F)** Heatmaps illustrating the expression profiles of the manually annotated fibroblast **(D)** or macrophage **(F)** subpopulations, respectively. UMAP, uniform manifold approximation and projection; αSMA, α-smooth muscle actin; APOE, apolipoprotein E; DCN, decorin; Fb, fibroblast; GSN, gelsolin; MΦ, macrophage; POSTN, periostin; SPP1, secreted phosphoprotein 1; vWF, von Willebrand factor.

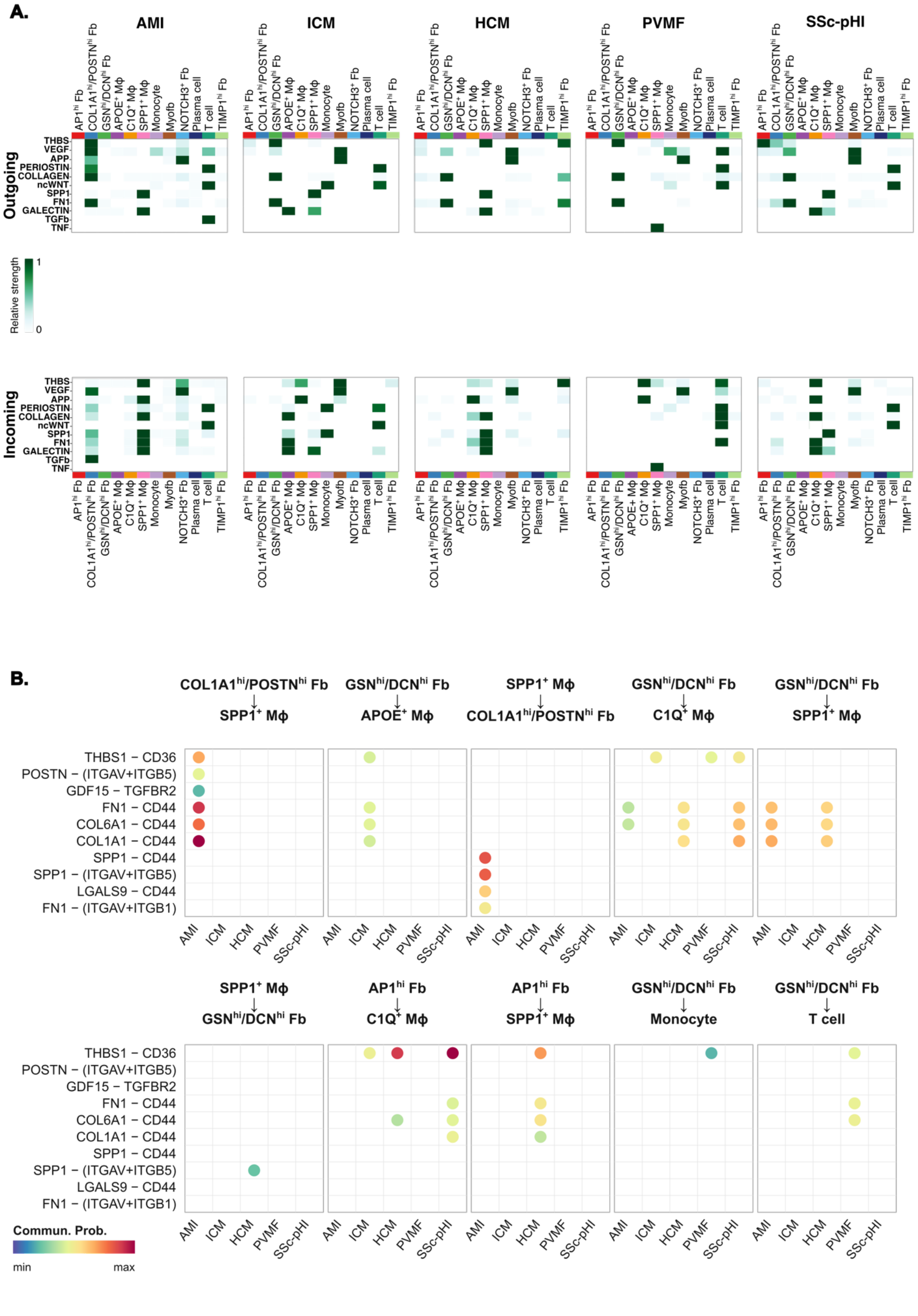
A.
AMI
ICM
HCM
PVMF
SSc-pHI
Outgoing
Incoming
THBS
VEGF
APP
PERIOSTIN
COLLAGEN
ncWNT
SPP1
FN1
GALECTIN
TGFb
TNF
AP1hi Fb
COL1A1hi/POSTNhi Fb
GSNhi/DCNhi Fb
APOE+ Mϕ
C1Q+ Mϕ
SPP1+ Mϕ
Monocyte
Myofb
NOTCH3+ Fb
Plasma cell
T cell
TIMP1hi Fb
Relative strength
1
0
B.
COL1A1hi/POSTNhi Fb → SPP1+ Mϕ
GSNhi/DCNhi Fb → APOE+ Mϕ
SPP1+ Mϕ → COL1A1hi/POSTNhi Fb
GSNhi/DCNhi Fb → C1Q+ Mϕ
GSNhi/DCNhi Fb → SPP1+ Mϕ
SPP1+ Mϕ → GSNhi/DCNhi Fb
AP1hi Fb → C1Q+ Mϕ
AP1hi Fb → SPP1+ Mϕ
GSNhi/DCNhi Fb → Monocyte
GSNhi/DCNhi Fb → T cell
THBS1 − CD36
POSTN − (ITGAV+ITGB5)
GDF15 − TGFBR2
FN1 − CD44
COL6A1 − CD44
COL1A1 − CD44
SPP1 − CD44
SPP1 − (ITGAV+ITGB5)
LGALS9 − CD44
FN1 − (ITGAV+ITGB1)
AMI
ICM
HCM
PVMF
SSc-pHI
Commun. Prob.
min
max

**Supplementary Figure 8: Cell-cell communication networks inferred by CellChat across disease conditions. (A)** Outgoing and incoming signaling patterns identified by CellChat for AMI, ICM, HCM, PVMF, and SSc-pHI. **(B)** Bubble plots showing selected significant fibroblast-immune ligand-receptor interactions across disease conditions, generated with CellChat's framework. "Commun. Prob." indicates communication probability from low to high as inferred by CellChat.

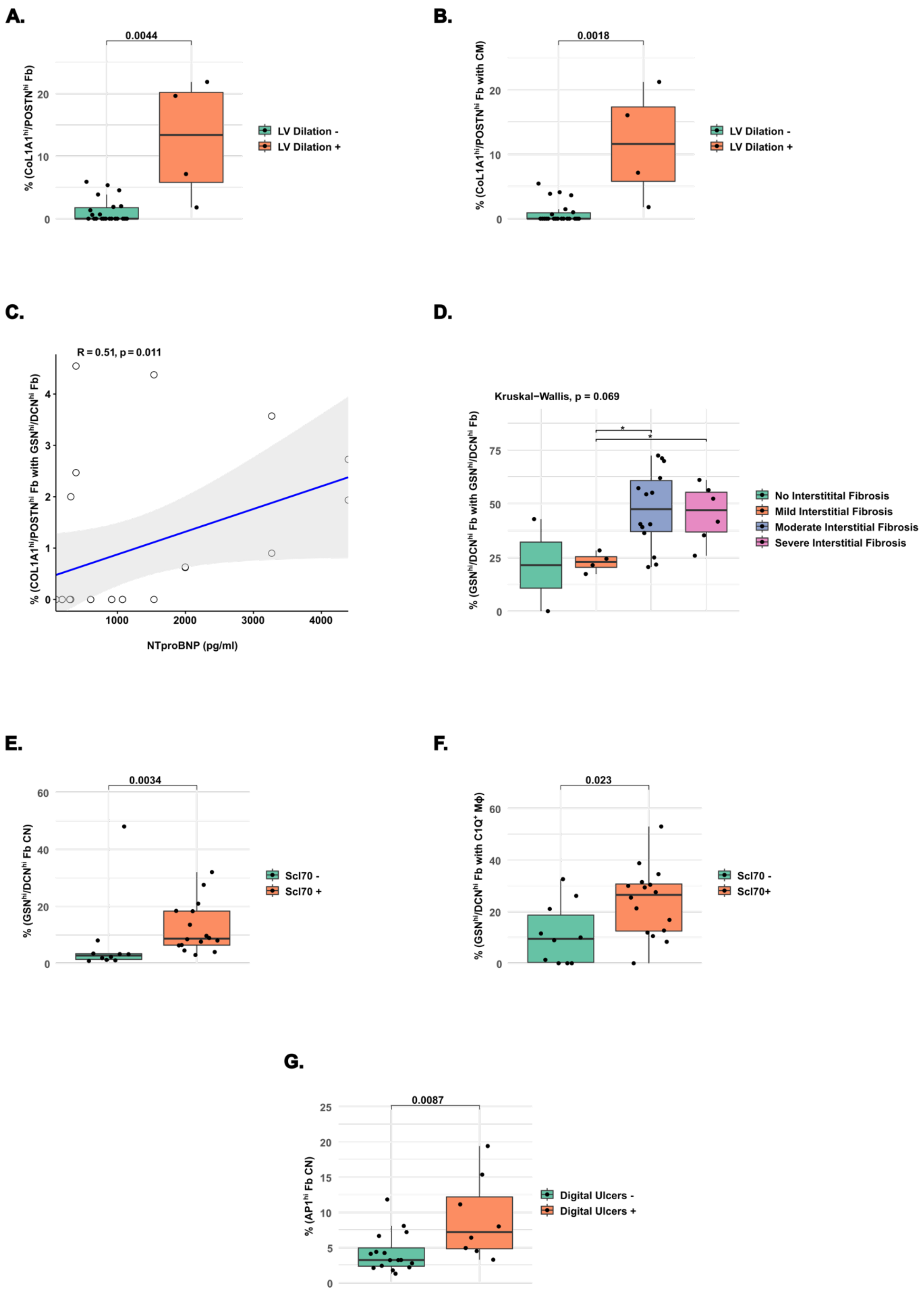
A.
0.0044
% (CoL1A1hi/POSTNhi Fb)
LV Dilation -
LV Dilation +
B.
0.0018
% (CoL1A1hi/POSTNhi Fb with CM)
LV Dilation -
LV Dilation +
C.
R = 0.51, p = 0.011
% (COL1A1hi/POSTNhi Fb with GSNhi/DCNhi Fb)
NTproBNP (pg/ml)
D.
Kruskal−Wallis, p = 0.069
% (GSNhi/DCNhi Fb with GSNhi/DCNhi Fb)
No Interstitital Fibrosis
Mild Interstitial Fibrosis
Moderate Interstitial Fibrosis
Severe Interstitial Fibrosis
E.
0.0034
% (GSNhi/DCNhi Fb CN)
Scl70 -
Scl70 +
F.
0.023
% (GSNhi/DCNhi Fb with C1Q+ MΦ)
Scl70 -
Scl70+
G.
0.0087
% (AP1hi Fb CN)
Digital Ulcers -
Digital Ulcers +

**Supplementary Figure 9: Association of fibroblast populations and neighborhoods with clinical characteristics in SSc-pHI patients. (A)** Box plot showing differences in the relative abundance of $COL1A1^{hi}/POSTN^{hi}$ fibroblasts in SSc-pHI patients with and without left ventricular (LV) dilation. **(B)** Box plot showing differences in the relative abundance of interactions between $COL1A1^{hi}/POSTN^{hi}$ fibroblasts and cardiomyocytes in SSc-pHI patients with and without LV dilation. **(C)** Scatter plot showing Spearman correlation analysis between the abundance of interactions between $COL1A1^{hi}/POSTN^{hi}$ and $GSN^{hi}/DCN^{hi}$ fibroblasts and NTproBNP levels. Each dot represents one sample. **(D)** Box plot showing the relative abundance of homotypic $GSN^{hi}/DCN^{hi}$ fibroblast interactions in SSc-pHI patients according to the histopathological severity of interstitial myocardial fibrosis, including patients without fibrosis. **(E)** Box plot showing differences in the relative abundance of $GSN^{hi}/DCN^{hi}$ fibroblast CN in SSc-pHI patients with and without anti-Scl70 autoantibodies. **(F)** Box plot showing differences in the relative abundance of interactions between $GSN^{hi}/DCN^{hi}$ fibroblasts and C1Q+ macrophages in SSc-pHI patients with and without anti-Scl70 autoantibodies. **(G)** Box plot showing differences in the relative abundance of $AP1^{hi}$ fibroblast niche in SSc-pHI patients with and without digital ulcers (DU). Statistical significance was assessed using Wilcoxon rank-sum tests or Kruskal-Wallis tests. Exact p values are shown. SSc-pHI, systemic sclerosis primary heart involvement; NTproBNP, N-terminal pro Brain natriuretic peptide.
